# Cortical response to naturalistic stimuli is largely predictable with deep neural networks

**DOI:** 10.1101/2020.09.11.293878

**Authors:** Meenakshi Khosla, Gia H. Ngo, Keith Jamison, Amy Kuceyeski, Mert R. Sabuncu

## Abstract

Naturalistic stimuli, such as movies, activate a substantial portion of the human brain, invoking a response shared across individuals. Encoding models that predict the neural response to a given stimulus can be very useful for studying brain function. However, existing neural encoding models focus on limited aspects of naturalistic stimuli, ignoring the complex and dynamic interactions of modalities in this inherently context-rich paradigm. Using movie watching data from the Human Connectome Project (HCP, *N* = 158) database, we build group-level models of neural activity that incorporate several inductive biases about information processing in the brain, including hierarchical processing, assimilation over longer timescales and multi-sensory auditory-visual interactions. We demonstrate how incorporating this joint information leads to remarkable prediction performance across large areas of the cortex, well beyond the visual and auditory cortices into multi-sensory sites and frontal cortex. Furthermore, we illustrate that encoding models learn high-level concepts that generalize remarkably well to alternate task-bound paradigms. Taken together, our findings underscore the potential of neural encoding models as a powerful tool for studying brain function in ecologically valid conditions.

## Introduction

How are dynamic signals from multiple senses integrated in our minds to generate a coherent percept of the world? Understanding the neural basis of perception has been a longstanding goal of neuroscience. Previously, sensory perception in humans has been dominantly studied via controlled task-based paradigms that reduce computations underlying brain function into simpler, isolated components, preventing broad generalizations to new environments or tasks (*1*). Alternatively, fMRI recordings from healthy subjects during free-viewing of movies present a powerful opportunity to build ecologically-sound and generalizable models of sensory systems, known as encoding models (*2, 3, 4, 5, 6, 7*).

To date, however, existing works on encoding models study sensory systems individually, and often ignore the temporal context of the sensory input. In reality, the different senses are not perceived in isolation; rather, they are closely entwined through a phenomenon now well-known as multi-sensory integration (*8, 9*). For example, specific visual scenes and auditory signals occur in conjunction and this synergy in auditory-visual information can enhance perception in animals, improving object recognition and event detection as well as markedly reducing reaction times (*10*). Furthermore, our cognitive experiences unfold over time; much of the meaning we infer is from stimulation sequences rather than from instantaneous visual or auditory stimuli. This integration of information from multiple natural sensory signals over time is crucial to our cognitive experience. Yet, previous encoding methodologies have precluded the joint encoding of this rich information into a mental representation of the world.

Accurate group-level predictive models of whole-brain neural activity can be invaluable to the field of sensory neuroscience. These models learn to disregard the idiosyncratic signals and/or noise within each individual, while capturing only the shared response relevant to the stimuli. Naturalistic viewing engages multiple brain systems and involves several cognitive processes simultaneously, including auditory and visual processing, memory encoding and many other functions (*11*). Group-level analysis in this paradigm is enabled by the synchrony of neuronal fluctuations in large areas of the cortex across subjects (*12*). Thus far, inter-subject correlation (ISC) analysis (*12*) has been a cornerstone tool for naturalistic paradigms because of its ability to characterize the shared response across individuals. Group-level encoding models adopt an alternative approach for capturing shared response, one grounded in out-of-sample prediction and generalization (*1*). This allows them to model neural activity beyond a constrained stimulus set. However, there is a clear gap between the two mediums of analysis. While ISC analysis suggests that large areas of the cortex exhibit fluctuations that are consistent across subjects, existing neural encoding models have largely focused on predicting activity within pre-defined functional areas of the brain such as visual and auditory cortices. It is unclear how they may be scaled to develop a single predictive model for whole-brain neural responses, given that naturalistic scenes produce wide-spread cortical activations. In this paper, we aim to fill this gap: provided adequate characterization of stimuli, we hypothesize that the stable component of neural activity across a subject population, i.e., the stimulus related activity, should be predictable. In the present study, we aim to quantify and improve the encoding of this wide-spread stimulus-driven cortical activity using rich stimulus descriptions.

Brain responses in real-world conditions are highly complex and variable. Owing to their high expressive capacity, deep neural networks (DNNs) are well-suited to model the complex high-dimensional nature of neural activity in response to the multitude of signals encountered during movie-watching. Recently, DNNs optimized for image or sound recognition have emerged as powerful models of computations underlying sensory processing (*4, 5, 7, 2*), surpassing traditional models of image or sound representation based on Gabor filters (*3*) and spectrotemporal filters (*13*), respectively, in higher-order processing regions. In this approach, the stimuli presented during brain activity recordings are fed as input to pre-trained neural networks and activations of individual layers are linearly transformed into predictions of neural responses in different regions of the brain. This approach affords a useful interpretation of these feature spaces as outcomes of a task-constrained optimization, shedding light on how high-level behavioral goals, such as recognition, may constrain representations in neural systems (*2*). While useful, task-driven features may diverge from optimal neural representations and tuning these features to better match the latter may be both feasible and beneficial (*14*). This approach can help bridge the quantitative gap in explaining neural responses under realistic conditions while improving our understanding of the nature of information processing in the brain. From a purely modeling standpoint, our methodological innovations are threefold. First, we propose an end-to-end deep-learning based encoding model that extracts semantic feature maps from audio and visual recognition networks and refines them jointly to predict the evoked brain response. To this effect, we demonstrate that using different modalities concurrently leads to improvements in brain encoding. Second, we note that cognitive perception during movie-watching involves maintaining memory over time and demonstrate the suitability of recurrent neural networks (RNNs) to capture these temporal dynamics. Finally, based on existing evidence of hierarchical information processing in visual and auditory cortices (*5, 7*), we adopt features at multiple levels of abstraction rather than low level or high level stimulus characteristics alone. We embed these inductive biases about hierarchy, long-term memory and multi-modal integration into our neural architecture and demonstrate that this comprehensive deep-learning framework generalizes remarkably well to unseen data. Specifically, using fMRI recordings from a large cohort of subjects in the HCP, we build group-level encoding models that reliably predict stimuli-induced neuronal fluctuations across large parts of the cortex. As a demonstration of application, we employ these encoding models to predict neural activity in response to other task-based stimuli and report excellent transferability of these models to artificial stimuli from constrained cognitive paradigms. This further suggests that these encoding models are able to capture high-level mechanisms of sensory processing.

Approaching multi-sensory perception through the predictive lens of encoding models has several advantages. Because of their unconstrained nature, encoding models can enable data-driven exploration and catalyze new discoveries. Using six neural encoding models with different temporal scales and/or sensory inputs, trained only on *∼*36 minutes of naturalistic data per subject, we can replicate findings from a large number of prior studies on sensory processing. First, by prominently highlighting the transition from short to long temporal receptive windows as we move progressively from early to high-level auditory areas, we can distinguish the cortical temporal hierarchy. Next, by differentiating uni-sensory cortices from multi-sensory regions such as the superior temporal sulcus and angular gyrus, we can reproduce the multi-modal architecture of the brain. Finally, by synthesizing neural response to arbitrary stimuli such as faces, scenes or speech, we can demonstrate the functional specialization of known brain regions for processing of these distinct categories. Altogether, our results highlight the advantages and ubiquitous applications of DNN encoding models of naturalistic stimuli.

## Materials and Methods

### Dataset

We study high-resolution 7T fMRI data of 158 individuals from the Human Connectome Project movie-watching protocol comprising 4 audio-visual movie scans (*15, 16*). The movies represent a diverse collection, ranging from short snippets of Hollywood movies to independent vimeo clips. All fMRI data was preprocessed following the HCP pipeline, which includes motion and distortion correction, high-pass filtering, head motion effect regression using Friston 24-parameter model, automatic removal of artifactual timeseries identified with Independent Component Analysis (ICA) as well as nonlinear registration to the MNI template space (*16*). Complete data acquisition and preprocessing details are described elsewhere (*15, 16*). Finally, whole-brain fMRI volumes of size 113×136×113 are used as the prediction target of all proposed encoding models. Rest periods as well as the first 20 seconds of every movie segment were discarded from all analysis, leaving ∼12 minutes of audio-visual stimulation data per movie paired with the corresponding fMRI response. We estimated a hemodynamic delay of 4 *sec* using ROI-based based encoding models, as the response latency that yields highest encoding performance (Figure S2, see Supplementary Information for details). Thus, all proposed models are trained to use the above stimuli to predict the fMRI response 4 seconds *after* the corresponding stimulus presentation. We train and validate our models on 3 audio-visual movies with a 9:1 split respectively and evaluate our models on the first three clips of the held-out test movie. Since the last clip in the held-out movie is repeated within the training movies, we excluded it from our analysis.

### Methodology

We train six encoding models employing different facets of the complex, dynamic movie stimulus. These include: (1) Audio-1sec and (2) Audio-20sec models, which are trained on single audio spectrograms extracted over 1 second epochs and contiguous sequences of 20 spectrograms spanning 20 seconds respectively; (3) Visual-1sec and (4) Visual-20sec models, trained with last frames of 1-second epochs and sequences of 20 evenly spaced frames within 20-second clips respectively; (5) Audiovisual-1sec and (6) Audiovisual-20sec models, which employ audio and visual input as described above, *jointly*. All models are trained to minimize the *mean squared error* between the predicted and measured whole-brain response. Figure 1 depicts the overall methodology for training different encoding models.

**Fig. 1.**
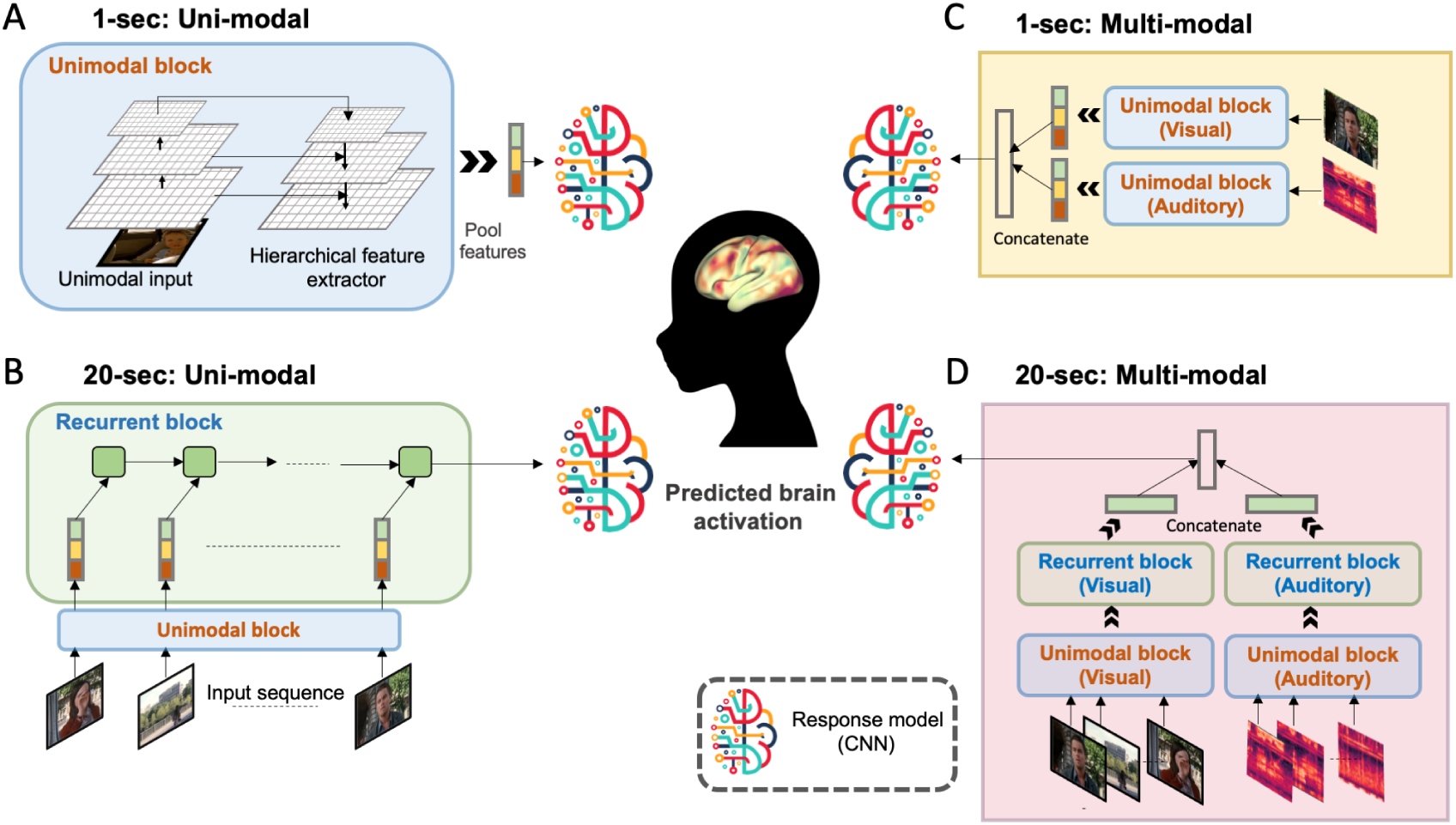
Schematic of the proposed models. (A) The short-duration (1-sec) auditory and visual models take a single image or spectrogram as input, extract multi-scale hierarchical features and feed them into a CNN-based response model to predict the whole-brain response (B) The long-duration (20-sec) uni-modal models take a sequence of images or spectrograms as input, feed their hierarchical features into a recurrent pathway and extract the last hidden state representation for the response model (C) The short-duration multi-modal model combines uni-modal features and passes them into the response model (D) The long-duration multi-modal model combines auditory and visual representations from the recurrent pathways for whole-brain prediction. Architectural details, including the feature extractor and convolutional response model are provided in Supplementary Information.

#### Stimuli

##### Audio

We extract mel-spectrograms over 64 frequency bands between 125-7500 Hz from sound waveforms to represent auditory stimulus in ∼1 second epochs, following (*17*). The audio spectrogram is treated as a single grayscale 96×64 image, denoted by 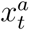, for the short duration model. For the longer-duration model, the input is simply a contiguous sequence of 20 of these gray-scale images, represented as 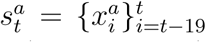. This representation of auditory input is also supported by strong evidence that suggests the cochlea may be providing a spectrogram-like input to the brain for information processing (*18*).

##### Visual

All videos were collected at 24 fps. We extract the last frame of every second of the video as a 720×1280×3 RGB input, denoted by 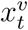, for the 1-sec models. We emphasize that the input here is a single RGB frame and we are using the 1-sec terminology only to be consistent with the nomenclature for audio models. We further arrange the last frame of every second in a 20-second clip into a sequence of 20 images, denoted by 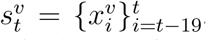, to represent the continuous stream of visual stimuli. These are presented to the longer-duration Visual-20sec and Audiovisual-20sec models.

The inputs to the Audio-1sec, Visual-1sec, Audio-20sec, Visual-20sec, Audiovisual-1sec and Audiovisual-20sec models are thus given as 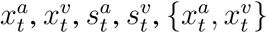 and 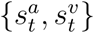 respectively.

#### Audio-1sec and Visual-1sec models

Neural encoding models comprise two components: a feature extractor, which pulls out relevant features, **s**, from raw images or audio waveforms and a response model, which maps these stimuli features onto brain responses. In contrast to existing works that employ a linear response model (*4, 7*), we propose a CNN-based response model where stimulus features are mapped onto neural data using non-linear transformations. Previous studies have reported a cortical processing hierarchy where low-level features from early layers of a CNN-based feature extractor best predict responses in early sensory areas while semantically-rich deeper layers best predict higher sensory regions (*7, 5*). To account for this effect, we employ a hierarchical feature extractor based on feature pyramid networks (*19*) that combines features from early, intermediate and later layers simultaneously. The detailed architectures of both components, including the feature extractor and convolutional response model are described in Figure S3. We employ state-of-the-art pre-trained ResNet-50 (*20*) and VGG-ish (*17*) architectures in the pyramid network to extract multi-scale features from images and audio spectrograms, respectively. The base architectures were selected because pre-trained weights of these networks optimized for behaviorally relevant tasks (recognition) on large datasets, namely Imagenet (*21*) and Youtube-8M (*22*), were publicly available. Resnet-50 was trained on image classification with 1000 classes, while the VGG-ish network was pre-trained on audio event recognition with ∼30K categories. Further, due to computational and memory budget, the Resnet-50 was frozen during training across all models. On the other hand, we were able to fine-tune the VGG-ish network in both the Audio and Audiovisual encoding models. We note that in contrast to images, there is a clear asymmetry in the axes of a spectrogram, where the distinct meanings of time and frequency might warrant 1D convolutions over time instead of 2D convolutions over both frequency and temporal axes. However, we found the benefits of a pre-trained network to be substantial in training convergence time and hence did not explore more appropriate architectures.

#### Audio-20sec and Visual-20sec models

Audio-20sec and Visual-20sec models employ the same feature extractor and CNN response model as their 1-second counterparts. However, here, the feature extraction step is applied on each image in a sequence of 20 frames, followed by a long short-term memory (LSTM) module to model the temporal propagation of these features. The output dimensions of the LSTM unit are set to 1024 and 512 for the visual and auditory models respectively, to ensure an equitable comparison with the corresponding 1-sec models. The last hidden state output of this LSTM unit is fed into the CNN response model with the same architecture as the 1-sec models.

#### Audiovisual-1sec and Audiovisual-20sec models

Meaningful comparison across different models requires the control of as many design choices as possible. To ensure fair comparisons, the Audiovisual-1sec model employs the same feature extractors as the Visual-1sec and Audio-1sec models. The only difference, here, is that the corresponding 1024-D and 512-D feature representations are concatenated before presenting to the CNN response model and the concatenated features are passed into a bottleneck layer to reduce the final feature dimensionality to the maximum among audio and visual feature dimensions, i.e., 1024, so that the multi-modal model is not equipped with a higher-dimensional feature space than the maximum among uni-modal models. We note that the response model has the same architecture across all 6 proposed models. Similarly, the Audiovisual-20sec model employs the same feature extraction scheme as the Visual-20sec and Audio-20sec models, but fuses the last hidden state output of the respective LSTM units by simple concatenation followed by a dense layer to reduce feature dimensionality to 1024 before feeding it into the response model.

### Evaluation

We first evaluated the prediction accuracy of all models on the independent held-out movie by computing Pearson correlation coefficient (R) between the measured and predicted response at every voxel. Here, the ‘measured’ response refers to the group-averaged response across the same group of 158 subjects on which the models were trained. Comparison among these models enables us to tease apart the sensitivity of individual voxels to input timescales and different sensory stimuli. Voxel-level correlation coefficients between the predicted and measured responses were averaged to summarize the prediction accuracy of each model in relevant cortical areas (Figure 2B-F). For this region-level analysis, ROIs were derived with a comprehensive multi-modal parcellation of the human cortex (*23*), which was mapped onto the MNI-1.6 mm resolution template. We note that ROIs were employed only to interpret the results of the study and relate them to existing literature. We emphasize that all performance metrics reported henceforth are based on voxel-level correlations. It is important to note that prediction accuracy at every voxel is bounded by the proportion of non-stimulus related variance that reflects measurement noise or other factors. We thus also show the regional level performance of all models against the reliability (“noise ceiling”) of measured responses within those regions (Figure 3).

**Fig. 2.**
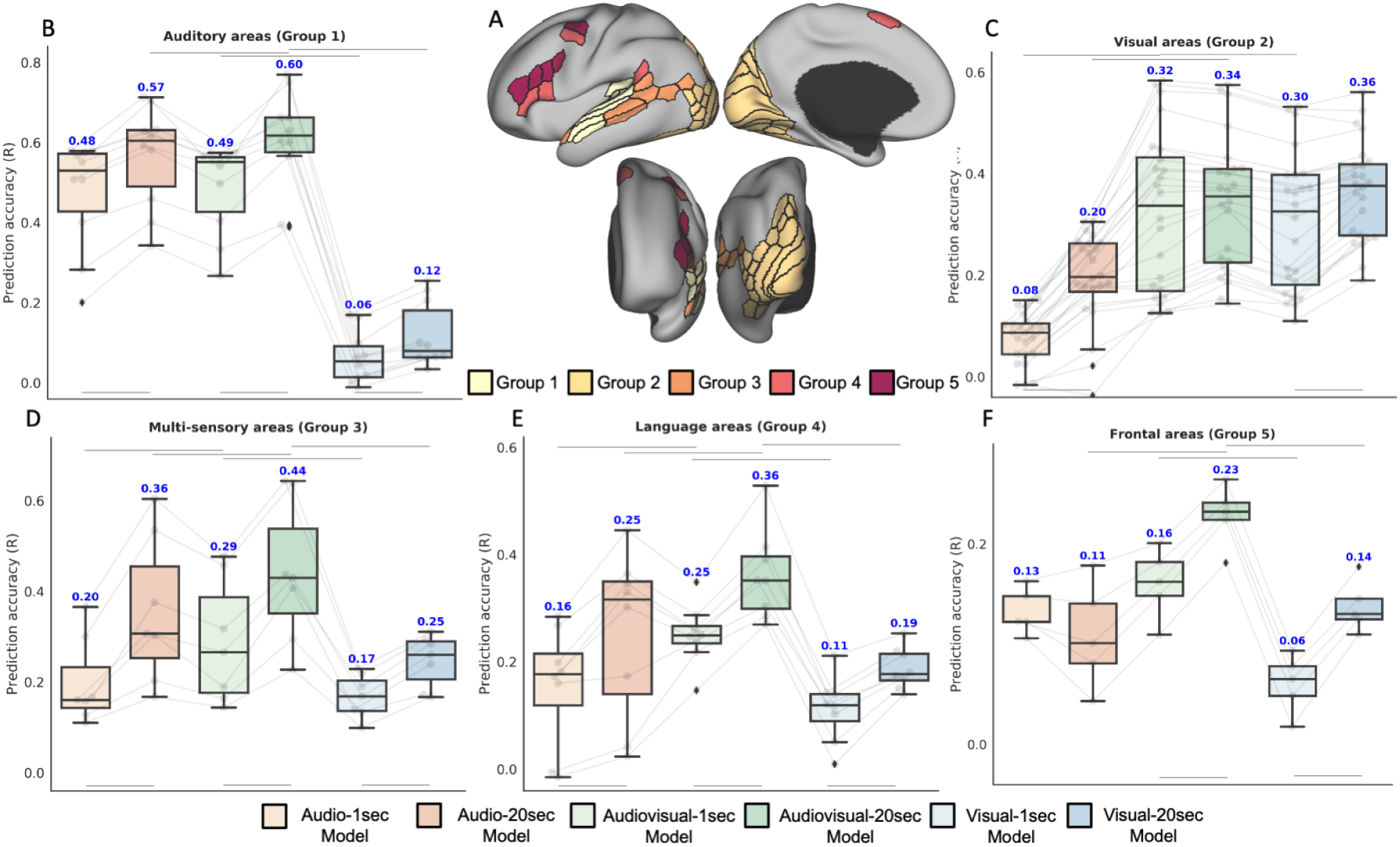
Regional predictive accuracy for the test movie. (B)-(F) depict quantitative evaluation metrics for all the proposed models across major groups of regions as identified in the HCP MMP parcellation (A). Predictive accuracy of all models is summarized across (B) auditory, (C) visual, (D) multi-sensory, (E) language and (F) frontal areas. Box plots depict quartiles and swarmplots depict mean prediction accuracy of every ROI in the group. For language areas (Group 4), left and right hemisphere ROIs are shown as separate points in the swarmplot because of marked differences in prediction accuracy. Statistical significance tests (results indicated with horizontal bars) are performed to compare 1-sec and 20-sec models of the same modality (3 comparisons) or uni-modal against multi-modal models of the same duration (4 comparisons) using paired t-test (p-value *<* 0.05, Bonferroni corrected) on mean prediction accuracy within ROIs of each group.

**Fig. 3.**
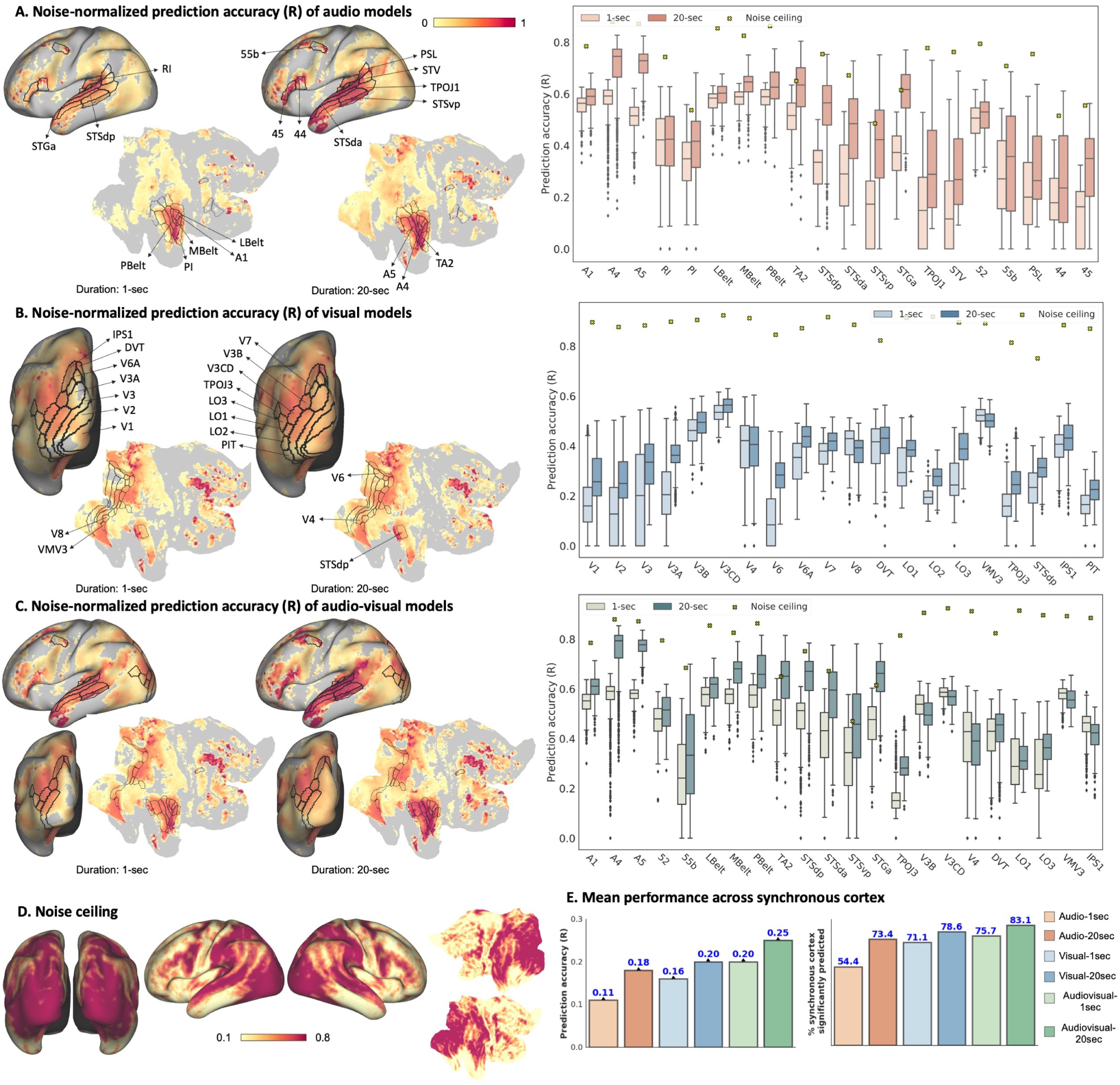
Predictive accuracy of uni-modal (A,B) and multi-modal (C) models over the whole brain in the test movie. Colors on the brain surface indicate the Pearson’s correlation coefficient between the predicted timeseries at each voxel and the true voxel’s timeseries normalized by the noise ceiling (D) computed on repeated validation clips. Only significantly predicted voxels (p-value *<* 0.5, FDR corrected) are colored. ROI box plots depict the un-normalized correlation coefficients between the predicted and measured response of voxels in each ROI and the respective noise ceiling for the mean. (E) shows the percentage of voxels in stimulus-driven cortex that are significantly predicted by each model and mean prediction accuracy across the stimulus-driven cortex.

#### Noise ceiling estimation

The reliability of the group-averaged response at each voxel is estimated from a short 84 second clip that was repeatedly presented at the end of all movie sessions. We compute an effective upper bound on our performance metric, i.e., the correlation coefficient, as the correlation between the measured fMRI response (group-mean) during different runs. We repeat this process 6 times (choosing pairs from 4 repeat measurements) to get a mean noise ceiling estimate per voxel, as shown in Figure 3D. We divide the voxel-level prediction accuracy (R) by this noise ceiling to get noise-normalized prediction accuracy of all models in left panels of Figure 3A-C. We note that this noise ceiling is computed on the repeated video clip, which is distinct from the test movie on which the model performance metrics are computed. Direct comparison against this noise ceiling can be sub-optimal, especially if the properties of the group-averaged response vary drastically across the two stimulus conditions. We address this limitation during model evaluation against data from a held-out independent group of subjects by computing a more suitable upper bound, which is achievable by a group-level encoding model (Figure S7, see Supplementary Information for more details). As we demonstrate in the results (Figure S7, S8), the trend and spatial distribution of model performance against noise ceiling remains unchanged across the model evaluation and noise ceiling estimation method.

## Results

### Multi-sensory inputs and longer time-scales lead to the best encoding performance with significant correlations across a large proportion of the stimulus-driven cortex

To gain quantitative insight into the influence of temporal history and multi-sensory inputs on encoding performance across the brain, we computed the mean prediction accuracy in five groups of regions defined as per the HCP MMP parcellation (*23*), namely, (1) auditory regions comprising both early and association areas, (2) early visual and visual association regions, (3) known multi-sensory sites and regions forming a bridge between higher auditory and higher visual areas, (4) language-associated regions, and (5) frontal cortical areas. As our research concerns stimulus-driven processing, only ROIs belonging to the “stimulus-driven” cortex were included in the above groups (Table S2, see Supplementary Information for the definition of “stimulus-driven” cortex). Groups 1 and 2, which are associated with a single modality (auditory or visual) do not show any marked improvement from audio-visual multi-sensory inputs and are best predicted by features of their respective sensory stimulus (Figure 2B,C). The performance boost with multi-sensory inputs is more pronounced in groups 3, 4 and 5 which are not preferentially associated with a single modality, but are involved in higher-order processing of sensory stimuli (Figure 2D-F). Further, temporal history of the stimulus yields consistent improvement in prediction performance in almost all groups of regions, albeit to different extents. Improvements in groups 3, 4 and 5 agree well with the idea that higher-order sensory processing as well as cognitive and perceptual processes, such as attention and working memory, are hinged upon the history of sensory stimuli; therefore, accumulated information benefits response prediction in regions recruited for these functions. Further, both auditory and visual association cortices are known to contain regions that are responsive to sensory information accumulated over the order of seconds (*24*). This potentially explains the significant improvement observed for long-timescale encoding models compared to their short-timescale counterparts in these sensory cortices (Figure 4). Together, the Audiovisual-20sec model integrating audiovisual multi-sensory information over longer time-scales yields maximum prediction accuracy (R) and highest percentage (∼83 percent) of significantly predicted voxels across the stimulus-driven cortex (Figure 3E), suggesting that the Audiovisual-20sec model can adequately capture complementary features of each additional facet (multi-sensory stimuli / temporal information) of the sensory environment.

**Fig. 4.**
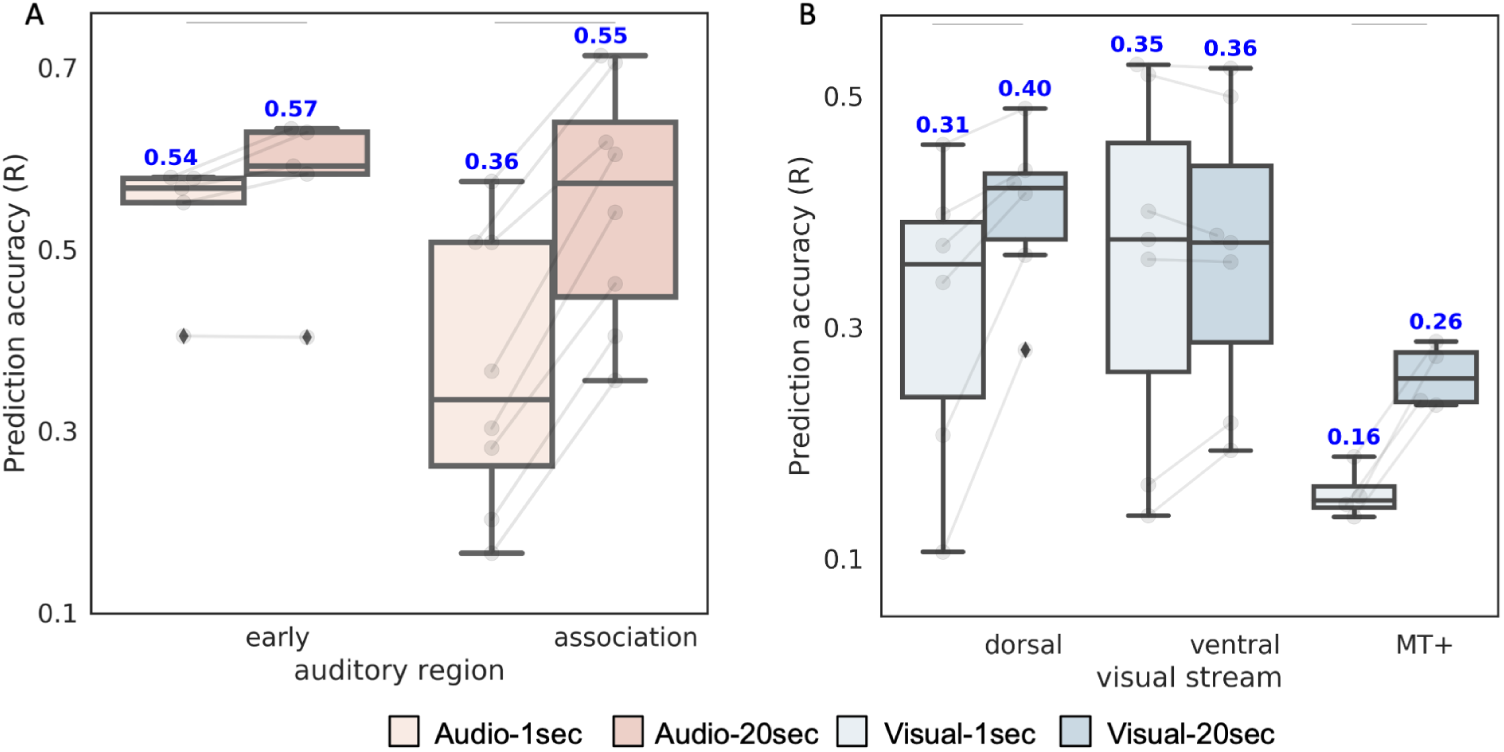
Influence of temporal history on encoding performance. (A) Mean predictive performance of Audio1sec and Audio-20sec models in early auditory and association auditory cortex ROIs. A major boost in encoding performance is seen across auditory association regions with the 20-sec model. (B) Mean predictive performance of Visual-1sec and Visual-20sec models across ROIs in the dorsal, ventral and MT+ regions. Dorsal stream and MT+ ROIs exhibit a significant improvement with Visual-20sec model but no effect is observed for the ventral stream. Boxplots are overlaid on top of the beeswarm plot to depict quartiles. Horizontal bars indicate significant differences between models in the mean prediction accuracy within ROIs of each stream using paired t-test (p-value *<* 0.05).

### Longer time-scales improve encoding performance, particularly in higher order auditory areas

As a movie unfolds over time, the dynamic stream of multi-modal stimuli continuously updates our neural codes. Evidence from neuroimaging experiments suggests that different brain regions integrate information at different timescales; a cortical temporal hierarchy is reported for auditory perception where early auditory areas encode short timescale events while higher association areas process information over longer spans (*25*). This temporal gradient of auditory processing is well-replicated within our study. Comparison of 1-sec and 20-sec models allows us to distinguish brain regions that process information at shorter timescales from those that rely on longer dynamics. There is a negligible contribution of longer timescale inputs on prediction correlations in regions within early auditory cortex, such as A1, LBelt, PBelt, MBelt and Restro-insular cortex (RI) (Figure 3A, 4A), in line with previous reports suggesting short temporal receptive windows (TRWs) of early sensory regions (*25*). Shorter integration windows are in agreement with the notion that these regions facilitate rapid processing of the instantaneous incoming auditory input. In contrast, response in voxels within auditory association ROIs lying mainly in the superior temporal sulcus or along the temporal gyrus (A4, A5, STSda, STSva, STSdp, STSvp, STGa, TA2) is seen to be much better predicted with longer time-scales (Figure 3A, 4A). Cumulatively across association ROIs, Audio-20sec model yields a highly significant improvement in prediction accuracy (*∼*50%) over the Audio-1sec model, in comparison to a marginal improvement (*∼*5%) across early auditory ROIs.

### Longer time-scales lead to significantly better predictions in the dorsal visual stream and MT+ complex

The distinct association of dorsal visual stream with spatial localization and action-oriented behaviors and ventral visual stream with object identification is well documented in the literature (*26*). Another specialized visual area is the medial temporal complex (MT+), which has been shown to play a central role in motion processing. The functional division between these streams thus suggests a stronger influence of temporal dynamics on responses along the dorsal pathway and MT+ regions. To test this hypothesis, we contrast the encoding performance of Visual-1sec and Visual-20sec models across the three groups by averaging voxel-wise correlations in their constituent ROIs. In accordance with the dorsal/ventral/MT+ stream definition in the HCP MMP parcellation, we use the following ROIs for analysis: (a) dorsal: V3A, V3B, V6, V6A, V7, IPS1 (b) ventral: V8, Ventral Visual Complex (VVC), PIT complex, Fusiform Face Complex (FFC) and Ventro-medial Visual areas 1,2 and 3 (c) MT+: MT, MST, V4t, FST. Figure 4B demonstrates the distribution of mean correlations over these ROIs for different models and streams. Our findings suggest that temporal history, as captured by the Visual-20sec model, can be remarkably beneficial to response prediction across the dorsal visual stream (30% improvement over Visual-1sec model) and the MT+ complex (62% improvement over Visual-1sec model), in agreement with our *a priori* hypothesis. Further, in our experiments, no marked improvement was observed for the ventral visual stream, indicating a non-significant influence of temporal dynamics on these regions.

### Auditory and visual stimuli features jointly approach the noise ceiling in multi-sensory areas

Examining prediction accuracy against response reliability allows us to quantify how far we are from explaining predictable neural activity. A high fraction of the stimulus-driven cortex (∼83%) is predictable with a longer timescale input and joint audiovisual features. Notably, areas extending anteriorly and posteriorly from the primary auditory cortex such as the posterior STS, STGa and TA2 achieve prediction correlations close to the noise ceiling with the Audiovisual20 sec model (Figure 3C), suggesting that DNN representations are remarkably suited to encode their response.

Interestingly, performance in auditory regions is much closer to the noise ceiling than visual regions. Understanding audition and vision in the same space further allows us to appreciate the differences between these modalities. While this may suggest that audition is perhaps a simpler modality to model, the differences could also result from a bias of the dataset. A more diverse sampling of acoustic stimuli in the training set could allow the model to generalize better in auditory regions. Furthermore, in contrast to auditory stimulation where all subjects hear the same sounds, visual stimulation can elicit highly varied responses dependent on gaze location. This variability could plausibly make group-level visual encoding a more difficult task.

### Joint encoding models tease apart the modal sensitivity of voxels throughout the sensory cortex

Neural patterns evoked by movies are not simply a conjunction of activations in modality-specific cortices by their respective uni-sensory inputs; rather, there are known cross-modal influences as well as regions that receive afferents from multiple senses (*27*). Can we interrogate a joint encoding model to reveal the individual contribution of auditory and visual features in encoding response across different brain regions? To address this question, we shuffled inputs of either modality along the temporal axis during inference. We measured test performance of the trained audio-visual model on predictions generated by shuffling inputs of one modality while keeping the other one intact. This distortion at test time allows us to identify areas that are preferentially associated with either visual or auditory modality. We hypothesized that regions encoding multi-sensory information will incur loss in prediction accuracy upon distortion of both auditory and visual information. Further, uni-sensory regions will likely be adversely affected by distortion of either auditory or visual information but not both. To test this hypothesis, we further developed a sensory-sensitivity index that directly reflects the sensitivity of individual brain regions to information about auditory or visual stimuli (see Supplementary Information for details). For this examination, we utilized the Audiovisual-1sec model to avoid potential confounds associated with temporal history, although analysis of the Audiovisual-20sec model showed similar results. Figure 5 demonstrates the result of this analysis on sensory-specific regions as well as regions known for their involvement in multi-sensory integration. The benefit from (non-distorted) multi-sensory inputs to the prediction correlations of the Audio-visual model is most remarkably seen in posterior STS, STGa and sensory-bridge regions such as the temporal-parietal-occipital junction (TPOJ1-3) and superior temporal visual (STV) area. Another region that seems to be employing features of both modalities, albeit to a lesser extent, is the frontal eye field (FEF), whose recruitment in audiovisual attention is well studied (*28*).

**Fig. 5.**
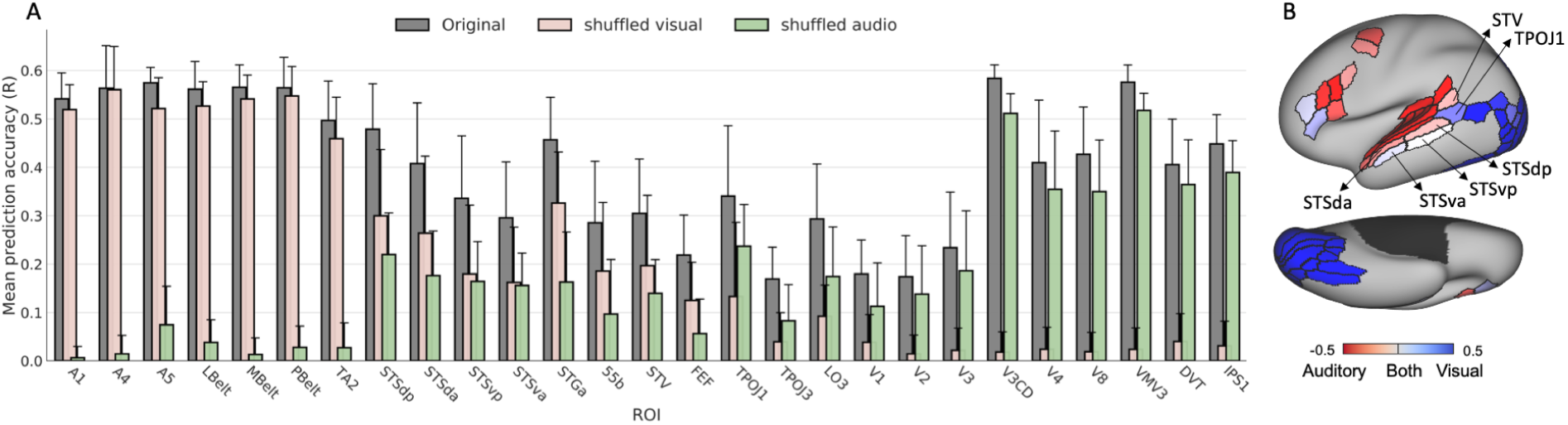
Sensitivity of ROIs to different sensory inputs. (A) Predictive accuracy (R) of audiovisual encoding model with and without input distortions, (B) Sensory sensitivity index of different brain regions as determined using performance metrics under input distortion (see Supplementary Information for details). Regions dominated by a single modality are shown in darker colors, whereas light-colored regions are better predicted by a combination of auditory and visual information. Red indicates auditory-dominant regions whereas blue indicates visual dominance.

Classically, multi-sensory integration hubs are identified as regions that show enhanced activity in response to multi-sensory stimulation as opposed to presentation of either uni-sensory stimuli based on some statistical criteria (*29*). Accordingly, the posterior STS is consistently described as a multi-sensory convergence site for audio-visual stimuli (*27, 30, 29, 9*). Its role in audiovisual linguistic integration has also been well-studied in the literature (*28*). Other multi-sensory integration sites reported extensively in prior literature include the temperoparietal junction (*9,27,28*) and superior temporal angular gyrus (*31*). Our findings above lend strong support for the multi-sensory nature of all these regions.

### Encoding models as virtual neural activity synthesizers

Next, we sought to characterize whether encoding models can generalize to novel task paradigms. By predicting neural activity for different visual categories from the category-specific representation task within the HCP Working Memory (WM) paradigm, we generated synthetic functional localizers for the two most common visual classes: faces and places. Specifically, we predict brain response to visual stimuli, comprising faces, places, tools and body parts, from the HCP task battery. We use the *predicted* response to synthesize contrasts (FACES-AVG and PLACES-AVG) by computing the difference between mean activations predicted for the category of interest (*faces* or *places* respectively) and the average mean activations of all categories at each voxel (Figure 6). The predicted and measured contrasts are thresholded to keep top 5% of the voxels.

**Fig. 6.**
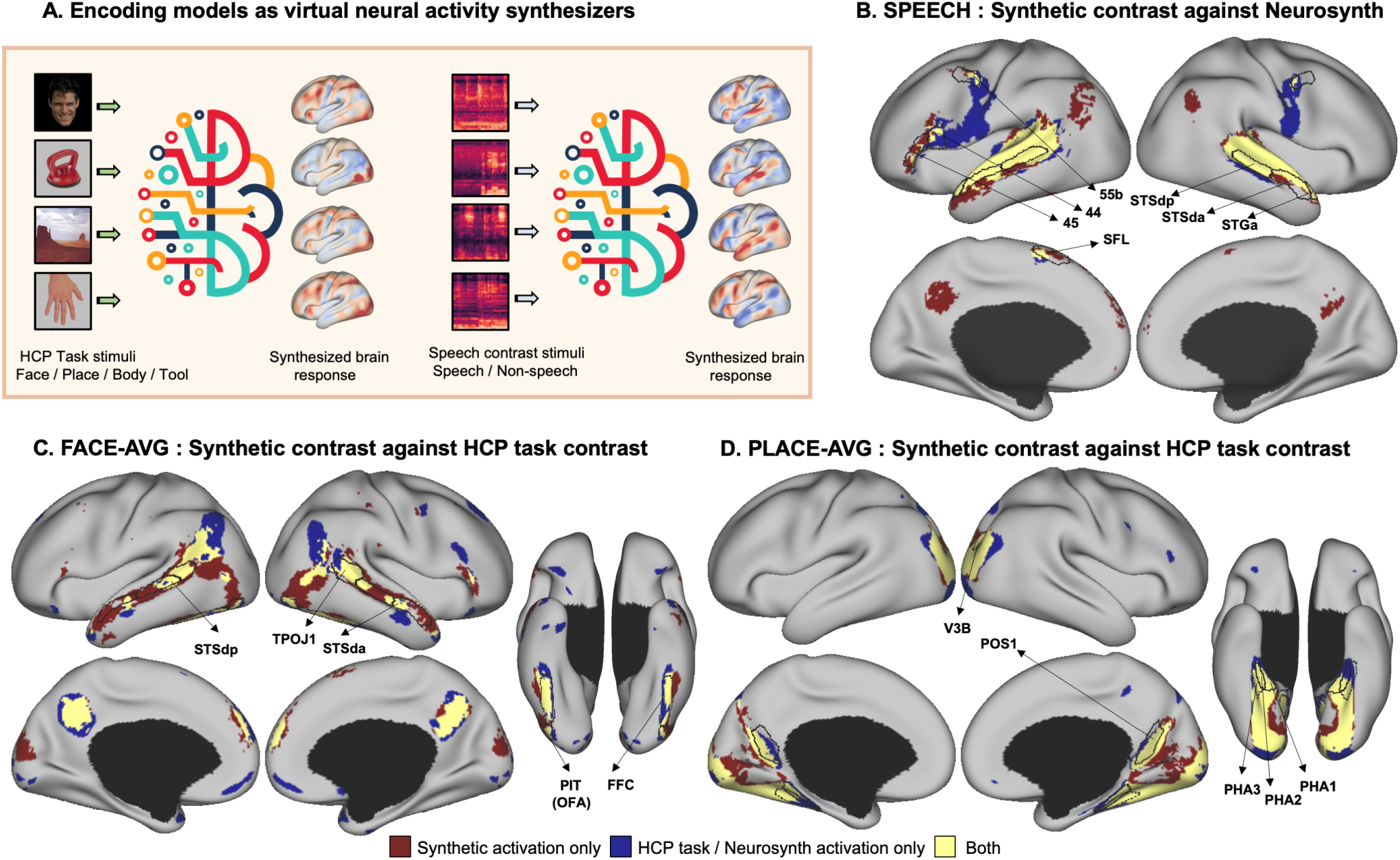
Encoding models as virtual brain activity synthesizers. (A) Synthetic contrasts are generated from trained encoding models by contrasting their “synthesized” (i.e., predicted) response to different stimulus types. (B) Comparison of the synthesized contrast for ‘speech’ against the speech association template on *neurosynth*. (C-D) compare the synthesized contrasts for ‘faces’ and ‘places’ against the corresponding contrasts derived from HCP tfMRI experiments.

We observe a notable overlap between the synthetic and measured group-level contrasts. Further, our findings are consistent with the well-known cortical specificity of neuronal activations for processing of *faces* and *places*. Both the synthetic and measured *faces* contrasts are consistent with previously identified regions for face-specific processing, including the fusiform face area (corresponds to fusiform face complex (FFC) in Figure 6), the occipital face area in lateral occipital cortex (overlaps with the PIT complex in HCP MMP parcellation), and regions within temporo-parieto-occipital junction and STS (*32, 33*). Among these, the *selective* role of the Fusiform Face Area in face processing has been most consistently and robustly established. Another region known to respond more strongly to faces than other object categories, namely posterior STS, has been previously implicated in processing of facial emotions (*32*).

Similarly, both synthetic and measured *places* contrasts highlight cortical regions thought to be prominent in selective processing of visual scenes. These include the parahippocampal areas (PHA1-3), retrosplenial cortex (POS1 in HCP MMP parcellation) and the transverse occipital sulcus (TOS), which comprises the occipital place area (OPA) (*34*).

Cortical areas related to speech processing are similarly discovered using our models by contrasting activations predicted for speech stimuli against non-speech stimuli such as environmental sounds (Figure 6B, see Supplementary Information for more details). The synthetic contrast shows increased activation in language-related areas of the HCP MMP parcellation such as 55b, 44 and the superior frontal language (SFL) area with left-lateralization, in accordance with previous language fMRI studies (*35*). In addition, areas tuned for voice processing in STS (*36*) are also highlighted. The synthetic map also shows highest correlation with ‘speech’ on *neurosynth* term-based meta-analysis (*37*) and overlaps considerably with the speech association template on the platform.

### Additional analyses

In prior studies, neural response prediction is done via regularized regression, where the signal at each voxel is modeled as a weighted sum of stimulus features with appropriate regularization on the regression weights. Following earlier works, we also train *l*_2_-regularized regression models using features derived from hierarchical convolutional networks trained on image or sound recognition such as those used in the proposed models, as well as semantic categories features labelled using the WordNet semantic taxonomy similar to (*38*). The latter are typically used for mapping the semantic tuning of individual voxels across the cortex. Our models consistently outperform the baselines, further illustrating the benefits of the proposed methodology (Figure S4(A)-(C), see Supplementary Information for more details). Additionally, we also performed ablation studies to understand the influence of different network components, namely the “non-linear” response model as well as the “hierarchical” feature extractor on model prediction performance and found that both components improve performance, although their relative contribution is stronger in visual encoding models than auditory models (Figure S4D, see Supplementary Information for more details). The superior predictive performance of our models in comparison to the classical approach along with our ablation studies suggest that an interplay of end-to-end optimization with a non-linear response model can jointly afford improved generalization performance.

To test the generality of the models beyond the subject population they were trained on, we further compared the predictions of all models against the group-averaged response of a held-out group within HCP comprising 20 novel subjects distinct from the 158 individuals used in the training set, on the same independent held-out movie. The noise ceiling for this group was computed as the correlation coefficient between the mean measured response for the *independent* test movie across all 158 subjects in the training set and the group-averaged response computed over the 20 new subjects. This metric captures the response component shared across independent groups of subjects and thus reflects the true upper bound achievable by a group-level encoding model. As shown in Figure S7 (see Supplementary Information for more details), the models can accurately predict neural responses as measured with respect to the group mean of the *held-out* subjects, with the Audiovisual-20sec model performance even approaching noise ceiling in some regions, particularly the higher-order auditory association regions and multisensory sites such as the posterior STS. Importantly, the predictivities across the cortical surface are consistent with the performance metrics reported for the training subject population in Figure 3. Finally, by comparing model predictions against neural responses at the single subject level for subjects from the held-out group, we further demonstrate that the Audiovisual-20sec model can also successfully capture the response component that individual subjects share with the population (Figure S9, see Supplementary Information for details).

## Discussion

Free viewing of dynamic audio-visual movies enables an ecologically valid analysis of a collective set of functional processes at once, including temporal assimilation and audio-visual integration in addition to momentary sensory-specific processing. Perception, under such stimulation, thus recruits sensory systems as well as areas subserving more sophisticated cognitive processing. Building quantitatively accurate models of neural response across widespread cortical regions to such real-life, continuous stimuli thus requires an integrated modelling of these disparate computations on sensory inputs. In this paper, we have presented six deep neural network based encoding models with varying sensory and temporal information about the audio-visual stimulus. Subsequently, we queried the role of input history and different sensory information on prediction performance across individual regions of the cortex. We have shown that exploiting the richness of the stimulus along the time axis and sensory modality substantially increases the predictive accuracy of neural responses throughout the cortex, so far as approaching the noise ceiling for voxels in some known multi-sensory sites, such as the posterior STS (*27, 30, 29, 9*).

Auditory and visual scenes are the principal input modalities to the brain during naturalistic viewing. Yet, existing encoding models ignore their interactions. We employ a common strategy in multi-modal machine learning settings, namely feature fusion, to jointly model auditory and visual signals from the environment. We find that minimizing the prediction error is a useful guiding principle to learn useful joint representations from an audio-visual stimulation sequence and demonstrate that models that consume multi-modal signals concurrently, namely, Audiovisual-1sec and Audiovisual-20sec, can not only predict the respective uni-modal cortices slightly better but also lead to remarkable improvements in predicting response of multi-sensory and frontal brain regions (Figure 2). Further, we show that multi-modal neural encoding models not only boost performance in large areas of the cortex relative to their uni-modal counterparts (Figure 2,3E), but also shed light on how neural resources are spatially distributed across the cortex for dynamic multi-sensory perception (Figure 5). The predictivity of different sensory inputs for neural response, as evaluated on independent held-out data, can facilitate reverse inference by identifying the sensory-associations of different brain regions, providing clues into the multi-sensory architecture of the cortex. By comparative analysis of predictive performance in different regions across models (Figure 2) as well as perturbation analysis within the multimodal model (Figure 5), we identify a number of regions that are consistently sensitive to both auditory and visual information, most notably the superior temporal sulcus and some frontal regions. Regions within inferior frontal cortex, have been implicated in the processing of visual speech, guiding sensory inferences about the likely common cause of multi-modal auditory and visual signals, as well as resolving sensory conflicts (*39*). Prior research has also implicated an extensive network of inferior frontal and premotor regions in comprehending audiovisual speech, suggesting that they bind information from both modalities (*40*). While unveiling the causal sequence of events for a mechanistic understanding of multi-sensory perception is not possible with the proposed approach, our findings align well with commonly held theories of sensory fusion which suggest that uni-sensory signals are initially processed in segregated regions and eventually fused in regions within superior temporal lobe, occipital-temporal junction and frontal areas (*27*). This proposition is corroborated by our experiments as response prediction in these regions is best achieved by a combination of both sensory inputs (Figure 3,5).

A linear response model with pre-trained and non-trainable feature extractors, while simple and interpretable, imposes a strong constraint on the feature-response relationship. The underlying assumption is that neural networks optimized for performance on behaviorally relevant tasks, are mappable to neural data with a linear transform. We designed a flexible model, capable of capturing complex non-linear transformations from stimulus feature space to neural space, leading to more quantitatively accurate models that are better aligned with sensory systems. Even better accounts of cortical responses are then obtained by interlacing dynamic, multimodal representation learning with whole-brain activation regression in an end-to-end fashion. Using these rich stimulus descriptions, we demonstrated a widespread predictability map across the cortex, that covers a large portion (∼83%) of the stimulus-driven cortex (Figure 3C,E), including association and some frontal regions. While inter-subject correlations in these regions are frequently reported (*12, 41*), suggesting their involvement in stimulus-driven processing, response predictability in these areas had remained elusive so far. Further, the cortical predictivity is maintained even as we compare model predictions against neural responses of held-out subjects (Figure S7 and S9), suggesting that the proposed models are capable of successfully capturing the “shared” or stimulus-driven response component. These results provide compelling evidence that deep neural networks trained end-to-end can learn to capture the complex computations underlying sensory perception of real-life, continuous stimuli.

We further demonstrated that encoding models can form an alternative framework for probing the time-scales of different brain regions. While primary auditory and auditory belt cortex (comprising A1, PBelt, LBelt, Mbelt) as well as the ventral visual stream benefit only marginally from temporal information, there is a remarkable improvement in prediction performance in auditory and visual association and pre-frontal cortices, most notably in superior temporal lobe, visuomotor regions within the dorsal stream such as V6A, temporal parietal occipital junction and inferior frontal regions. The improvement in prediction performance with the 20-second input is consistently seen for both uni-modal and multi-modal models. It is important to acknowledge that directly comparing the prediction accuracies of static (1-sec) and recurrent (20-sec) models to infer processing timescales of different brain regions has its limitations. First, this analysis can be confounded by the slow hemodynamic response as performance improvement may be driven in part by the slow and/or spatially varying dynamics. Based on our analysis with ROI-level encoding models, the latter seems like a less plausible explanation

(Figure S2, see Supplementary Information for details). Further, we performed additional analyses to understand the relationship between performance improvement in individual voxels and their autocorrelation properties and found a strong correspondence between the two, suggesting that the distribution of performance improvement across the cortex broadly agrees well with processing timescales (Figure S5, see Supplementary Information for details).

Predictions from long-timescale models are based on temporal history as provided in stimulus sequences, and not just the instantaneous input. Modeling dynamics within these sequences appropriately is crucial to probe effects of temporal accumulation. RNNs have internal memories that capture long-term temporal dependencies relevant for the prediction task, in this case, encoding brain response, while discarding task-irrelevant content. We compare this modeling choice against a regularized regression approach on stimulus features concatenated within T-second clips, with T ranging between 1 and 20 (Figure S4, see Supplementary Information for details). The inferior performance compared to our proposed models as well as a non-increasing performance trend against T for these linear models indicates that accumulation of temporal information by simply concatenating stimulus features over longer temporal windows is insufficient; rather, models that can efficiently store and access information over longer spans, such as RNNs with sophisticated gating mechanisms, are much more suitable for modeling neural computations that unfold over time. Since activations of units within RNNs depend not only on the incoming stimulus, but also on the “current” state of the network as influenced by past stimuli, they are capable of holding short-term events into memory. Adding the RNN module can thus be viewed as augmenting the encoding models with working memory.

Investigating timescales of representations across brain regions by understanding the influence of contextual representations on language processing in the brain, as captured by LSTM language models for instance, has become a major research focus recently (*42*). In these language encoding models for fMRI, past context has been shown to be beneficial in neural response prediction, surpassing word embedding models. However, models that explain neural responses under dynamic natural vision while exploiting the rich temporal context have not yet been rigorously explored with human fMRI datasets. In a previous study with awake mice, recurrent processing was shown to be useful in modelling the spiking activity of V1 neurons in response to natural videos (*43*). In dynamic continuous visual stimulation fMRI paradigms, a common practice is to concatenate multiple delayed copies of the stimulus to model the hemodynamic response function as a linear finite impulse response (FIR) function (*38*). However, since the feature dimensionality scales linearly with time-steps, this approach is limited to HRF modeling and is not feasible to capture longer dynamics of the order of tens of seconds. Another approach is to employ features from neural networks trained on video tasks, such as action recognition (*6*). However, these encoding models are constrained to capture one aspect of dynamic visual scenes and are likely useful to predict neural responses in highly localized brain regions. Most studies in visual encoding remain limited to static stimuli and evoked responses in relatively small cortical populations.

Our brain has evolved to process ‘natural’ images and sounds. In fact, recent evidence has shown that sensory systems are intrinsically more attuned to features of naturalistic stimuli and such stimuli can induce stronger neural responses than task-based stimuli (*44*). Here, we demonstrate that encoding models trained with naturalistic data are not limited to modeling responses of their constrained stimuli set. Instead, by learning high-level concepts of sensory processing, these models can also generalize to out-of-domain data and replicate results of alternate task-bound paradigms. While our models were trained on complex and cluttered movie scenes, we tested their ability to predict response to relatively simple stimuli from HCP task battery, such as faces and scenes (Figure 6). The remarkable similarity between the predicted and measured contrasts in all cases suggests that ‘synthetic’ brain voxels, predicted by the trained DNNs, correspond well with the target voxels they were trained to model. We thus provide evidence that these encoding models are capsulizing stimulus-to-brain relationships extending beyond the experimental world they were trained in. On the other hand, classical fMRI experiments, for instance task contrasts, don’t generalize outside the experimental circumstance they were based on. This preliminary evidence suggests that encoding models can serve as promising alternatives for circumventing the use of contrast conditions to study hypotheses regarding the functional specialization of different brain regions. Embedded knowledge within these descriptive models of the brain, could also be harnessed in other applications, such as independent neural population control by optimally synthesizing stimuli to elicit a desired neural activation pattern (*45*).

With purely data-driven exploration of fMRI recordings under a hypothesis-free naturalistic experiment, our models replicate the results of previous neuroimaging studies operating under controlled task-based regimes. Our analysis lends support to existing theories of perception which suggest that primary sensory cortices build representations at short timescales and lead up to multi-modal representations in posterior portions of STS (*25*). Encoding performance in these regions is consistently improved with longer timescales as well as multi-sensory information. We reasoned that regions that are sensitive to multi-modal signals and/or longer stimulus dynamics could be distinguished by interrogating the performance of these models on unseen data. To date, encoding models have been rarely used in this manner to assess integration timescales or sensory-sensitivity of different brain regions. Classically, processing timescales have been probed using various empirical strategies, for example, by observing activity decay over brief stimulus presentations or by comparing auto-correlation characteristics of resting-state and stimulus-evoked activity (*46*). Further, multi-sensory regions are identified via carefully-constructed experiments with uni-modal and multi-modal stimulus presentations, followed by analysis of interaction effects using statistical approaches (*27*). Here, we suggest that encoding models can form an alternate framework to reveal clues into these functional properties that can be rigorously validated with future investigation. As with interpreting the results of any predictive model, one should, however, proceed with caution. Sounds are generated by events; this implies that sound representations implicitly convey information about actions that generated them. Similarly, visual imagery provides clues into auditory characteristics, such as the presence of absence of speech. Thus, it is difficult to completely disentangle the individual contributions of auditory and visual features to prediction performance across cortical regions. Similarly, longer time-scale inputs can lead to a more robust estimate of the momentary sensory signal, potentially confounding the interpretations of TRWs. Here, we contend that these models can, nonetheless, serve as powerful hypothesis generation tools.

The methodological innovations in this study must also be considered in light of their limitations. Due to high dimensionality of features in early layers of the ResNet architecture for high-dimensional visual inputs, we employ pooling operations on these feature maps. Thus, low-level visual features, such as orientations, are compromised. The consequent unfavorable outcome is a low predictive performance in V1. Further, since different subjects can focus on different parts of the stimulus, group-level models can also blur out the precise object orientation information. This is particularly relevant for complex naturalistic stimuli such as movies. In the future, incorporating eye gaze data into these models can be an interesting exploration. Furthermore, due to computational constraints, the proposed model is only able to examine the effects of stimuli up to 20 seconds in the past. However, previous research with naturalistic stimuli has shown that some brain regions maintain memory of the order of minutes during naturalistic viewing (*47*). Existing evidence also suggests that neural activity is structured into semantically meaningful and coherent events (*25*). Capturing long-range context in encoding models can be a challenging, yet fruitful endeavour yielding potentially novel insights into memory formation.

There are also inherent differences between proposed neural network models and biological networks. DNNs fail to capture known properties of biological networks such as local recurrence, however, they have been found to be useful for modelling neural activity across different sensory systems. At present, feed-forward DNNs trained on recognition tasks constitute the best predictors of sensory cortical activations in both humans and non-human primates (*2*). In light of this observation, a recent study proposed that very deep feed-forward only CNNs (for example, ResNet-50 as employed in this study for visual feature extraction) might implicitly be approximating ‘unrolled’ versions of recurrent computations of the ventral visual stream (*48*). Object recognition studies on non-human primates have also hinted at a functional correspondence between recurrence and deep non-linear transformations (*49*). Although the functional significance of intra-regional recurrent circuits in core object recognition is still under debate, mounting evidence suggests they may be subserving recognition under challenging conditions (*49, 50*). Thus, investigation of more neurobiologically plausible models of the cortex that innately model intra-regional recurrent computations should be explored in the future, especially in relation to their role in visual recognition.

## Concluding remarks

Comprehensive descriptive models of the brain need comprehensive accounts of the stimulus. Using a novel group-level encoding framework, we showed that ‘reliable’ cortical responses to naturalistic stimuli can be accurately predicted across large areas of the cortex using multisensory information over longer time-scales. Since our models were trained on a large-scale, multi-subject and open-source dataset, we believe these results could provide an important point of reference against which encoding models for naturalistic stimuli can be assayed in the future. The continued interplay of artificial neural networks and neuroscience can pave the way for several exciting discoveries, bringing us one step closer to understanding the neural code of perception under realistic conditions.

## H2: Supplementary Materials

Supplementary Text

Figs. S1 to S9

Tables S1 to S2

References *(51-53)*

## Acknowledgments

This work was supported by NIH grants R01LM012719 (MS), R01AG053949 (MS), R21NS10463401 (AK), R01NS10264601A1 (AK), the NSF NeuroNex grant 1707312 (MS), the NSF CAREER 1748377 grant (MS) and Anna-Maria and Stephen Kellen Foundation Junior Faculty Fellowship (AK).

## Data and Software availability

All experiments in this study are based on the Human Connectome Project movie-watching database. The dataset is publicly available for download through the ConnectomeDB software (https://db.humanconnectome.org/). Throughout this study, we utilized 7T fMRI data from the ‘Movie Task fMRI 1.6mm/59k FIX-Denoised’ package within HCP. The network implementation, analysis codes as well as trained model weights will be made available on the project Github page.

## Supplementary materials

### HCP Movies

Table S1 summarizes the HCP movie-watching dataset split used for training and evaluating all models.

**Table S1.**
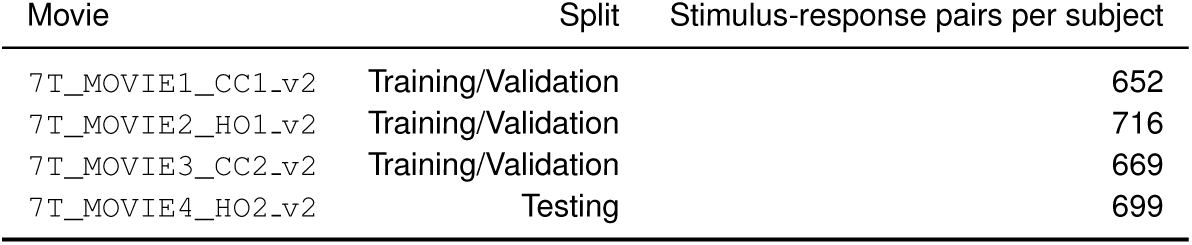
HCP dataset split

### Region of Interest (ROI) selection

ROIs were selected for each analysis based on the descriptions provided in the neuroanatomical supplementary results of the HCP MMP parcellation (*23*) and an extensive literature review. For Figure 2 in the main text and Figure S8, ROIs were thus assigned to groups 1-5 according to Table S2).

**Table S2.** ROI categorization

| Group | ROIs |
| --- | --- |
| 1. Auditory | A1, LBelt, PBelt, MBelt, RI, STSda, STSva, A4, A5, TA2 |
| 2. Visual | V1, V2, V3, V3A, V3B, V3CD, V4, V4t, V6, V6A, V7, V8, DVT, LO1-3, PIT, FFC, VMV1-3, IPS1, MT, VVC |
| 3. Multi-sensory + sensory bridges | STSdp, STSvp, STGa, STV, TPOJ1-3 |
| 4. Language | 55b, SFL, PSL, 44, 45 |
| 5. Frontal | IFSa, IFSp, IFJa, IFJp, FEF |

Dorsal and ventral visual stream ROIs as well as early and association auditory cortex ROIs in Figure 4 (main text) were derived from the explicit stream segregation and categorization described in the HCP MMP parcellation (*23*) and are defined here for quick reference.

- Dorsal: V3A, V3B, V6, V6A, V7, IPS1
- Ventral: V8, VVC, PIT, FFC, VMV1-3
- MT+: MT, MST, V4t, FST
- Early auditory: A1, PBelt, MBelt, RBelt, RI
- Association auditory: A4, A5, TA2, STGa, STSdp, STSda, STSvp, STSva

All ROIs are shown in Figure S1

**Fig. S1.**
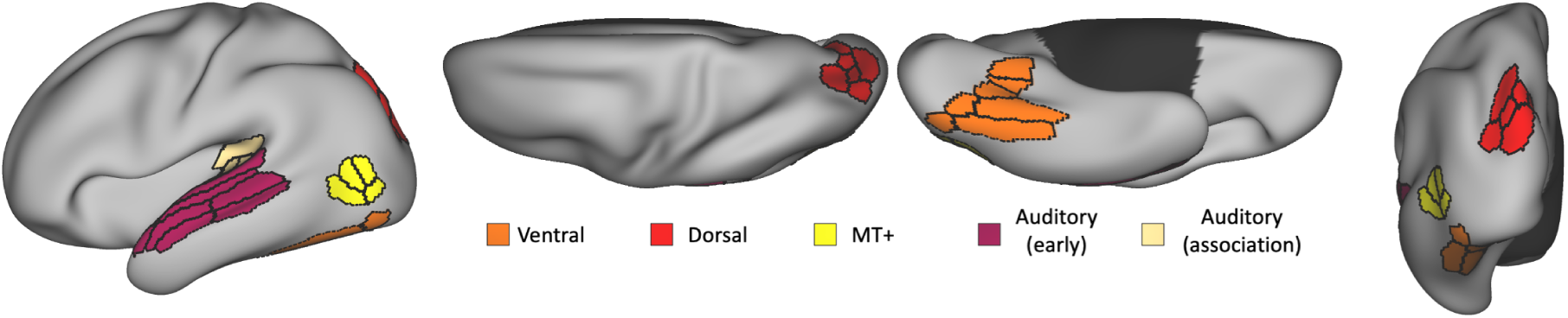
Group segregation from the HCP MMP parcellation.

### Estimating BOLD response delay

BOLD response delay was estimated using ROI-level encoding models due to their faster iteration times in comparison to voxel-wise encoding. The input to these models was the preprocessed stimuli as described for voxel-wise encoding with the same train-validation-test split, and the output was the evoked ROI-level fMRI response at different lags (1-7 seconds) from the stimulus. Thus, the output is a 360-D vector corresponding to the mean fMRI response in each ROI of the HCP MMP parcellation. The feature extractors were identical to those in the proposed voxel-wise auditory and visual models. However, instead of a convolutional response model, here, the response model comprised two fully connected layers with output dimensions of 512 and 360 with an exponential linear unit and linear activation respectively. All models were trained for 20 epochs with a batch size of 4 and a learning rate of 1e-4. Validation curves were monitored to ensure convergence. Prediction accuracy of each model was computed as the mean Pearson’s correlation coefficient between the predicted and measured response across all ROIs, in the held-out movie dataset. Based on Figure S2, we estimated a response delay of 4 seconds, as this lag yielded the maximum prediction accuracy across all ROIs for both auditory and visual ROI-level models. Further, even while restricting the prediction accuracy (R) to ROIs within different cortical areas (such as the early/association auditory areas or the dorsal/ventral visual stream), the optimal lag was consistently 4 seconds, suggesting that the difference in performance of 1-sec and 20-sec models in these regions (Figure 4) is not largely driven by differences in the hemodynamic response function (HRF).

**Fig. S2.**
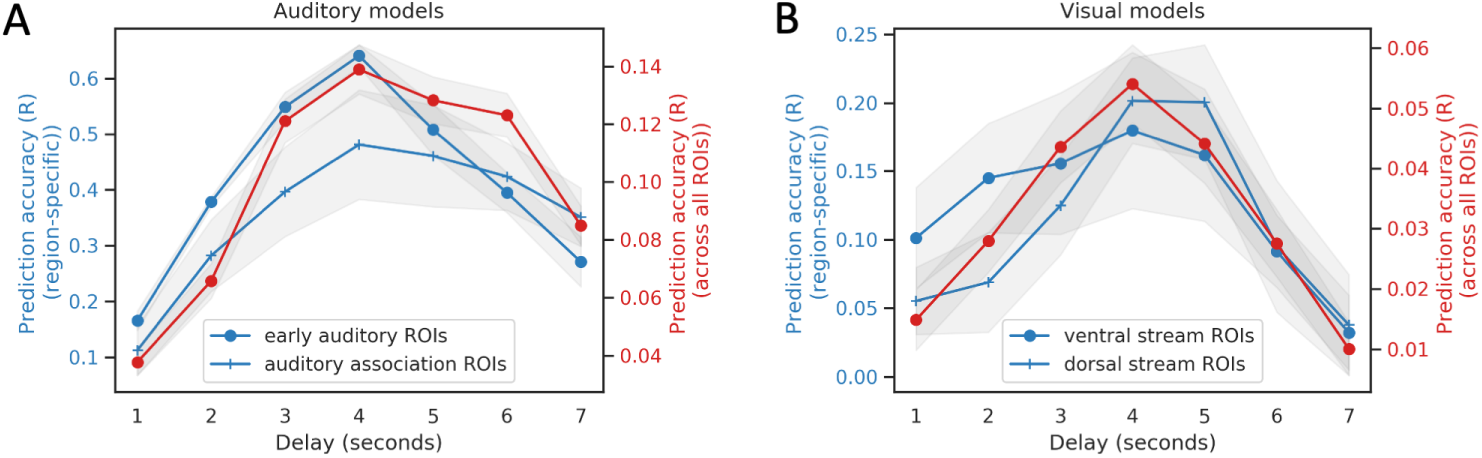
ROI-based encoding performance for estimating delay. (A) depicts the estimated mean and standard error of the prediction accuracy (R) across various delays (1-7s) within the early auditory and association auditory group (blue) as well as across all ROIs (red), as obtained using the single epoch (1s) auditory model. (B) depicts the estimated mean and standard error of the prediction accuracy (R) for various delays (1-7s) within the primary and dorsal visual streams (blue) as well as across all ROIs (red), as obtained using the single frame visual model. Gray regions depict the standard error in estimating mean across ROIs within each group. ROI categorization is described in the sub-section on ROI selection.

### Defining the stimulus-driven or “synchronous” cortex

We isolated voxels involved in stimulus-driven processing, termed “synchronous” or “stimulus-driven” voxels, by computing mean inter-group correlations over all training movies. Intergroup correlations were computed by splitting the entire group of subjects into two halves and computing correlations between the mean response time-course of each half (comprising 79 subjects) at every voxel. We employed a liberal threshold of 0.15 for this correlation value. Thus, the mask of “stimulus-driven” voxels included those voxels that achieved an inter-group correlation of 0.15 or above. We computed mean quantitative metrics over this mask in Figure 3E (main text) to compare different models.

### Model architectures and implementation

The base feature extraction networks and convolutional response model in Figure 1 had the architecture as detailed in Figure S3. The feature extraction networks are reminiscent of the feature pyramid network, which has shown significant improvements as a generic feature extractor across various applications. These networks comprise a parallel top-down pathway with lateral connections which grants them the ability to characterize both “what” and “where” in cluttered scenes, thereby enhancing object detection. We note that similar models with top-down and skip connections have been popular in vision research, since they can enrich low-level features with high-level semantics. The output of the feature extractor is fed into the convolutional response model to predict the evoked fMRI activation. This enables us to train both components of the network simultaneously in an end-to-end manner. Since the output response is differentiable with respect to network weights, the weights are adjusted via a first-order gradient-based optimization method to minimize the *mean squared error* between the predicted and target activation values across the entire brain.

For ResNet-50, we use activations of the last residual block of each stage, namely, res2, res3, res4 and res5 to construct our stimulus descriptions **s**. From the VGG-ish network, we use the activations of each convolutional block, namely, conv2, conv3, conv4 and the penultimate dense layer fc2 ^1^. The first three set of activations are refined through a top-down path to enhance their semantic content, while the last activation is concatenated into **s** directly (res4 activations are vectorized using global average pool). The top-down path comprises three feature maps at different resolutions with an up-sampling factor of 2 successively from the deepest layer of the bottom-up path. Each such feature map comprising 256/128 channels (in visual/auditory models respectively) is merged with the corresponding feature map in the bottom-up path (reduced to 256/128 channels by 1×1 convolutions) by element-wise addition. Subsequently, the feature map at each resolution is collapsed into a 256/128 dimensional feature vector through a global average pool operation and concatenated into **s**, leading to a 1024-D and 512-D feature representation for the visual and auditory stimuli respectively. The aggregated features are then passed onto a CNN comprising the following feedforward computations: a fully connected layer to map the features into a vector space which is reshaped into a 1024-channel cuboid of size 6×7×6 followed by four 3×3×3 transposed convolutions (conv.T) with a stride of 2 and exponential linear unit activation function to up-sample the latter. Each convolution reduces the channel count by half with the exception of the last convolution which outputs the single-channel predicted fMRI response.

The 20-second models additionally comprised an LSTM layer to model the temporal propagation of features across the contiguous sequence of input frames and/or spectrograms. The LSTM module has driven success across varied sequence modeling tasks due to its ability to efficiently regulate the flow of information across cells through gating. The memory cell in LSTM is modulated by three gates, namely, the input, forget and output gates. We note that the LSTM layer did not change the dimensionality of the input features so that equitable comparisons can be made against 1-sec models. The Audiovisual-1sec model concatenated features obtained from the base visual (1024-D) and audio (512-D) feature extraction networks, reduced their combined dimensionality to the higher value among the two (1024-D) by passing through a bottleneck dense layer followed by the same convolutional response model. The Audiovisual-20sec model additionally incorporated modal-specific LSTM networks prior to feature concatenation.

#### Implementation

We note that all 6 models have roughly the same order of trainable parameters in the range of 242M-362M. All parameters were optimized using Adam with a learning rate of 1e-4. Auditory and visual models were trained for 50 epochs with unit batch size. The stimulus as well as subject whose fMRI response is used as the target in the loss (“mean squared error”) are randomly sampled over each step of the training but kept consistent across models. We found this method to work better than using the group-averaged response as target, presumably because this sampling provides information about both the cross-subject mean and the variance of response. Given the noise characteristics at each voxel, we hypothesize that this enables the model to focus on regions that can be well predicted with the given stimulus. Validation curves were monitored for all models to ensure convergence.

### Regularized linear regression: WordNet features

Another popular approach in voxel-wise forward encoding beyond primary sensory cortices is the semantic category encoding model that is based on high-level semantic features (*38*). This approach relies on labels that indicate the presence of semantic object and action categories in each movie frame. In this analysis, we employed WordNet labels that were provided as part of the HCP movie-watching data pipeline. The semantic labels were manually assigned by the Gallant lab team using the WordNet semantic taxonomy and subsequently converted to WordNet synsets to build an 859-D semantic representational space (corresponding to 859 WordNet synset names). Following (*38*), we fitted *l*_2_ regularized linear regression models (known as ridge regression) to find weights corresponding to different input features for every voxel. The regularization parameter, *α* was optimized independently for each voxel by testing among 10 log-space values in [1, 1000]. The optimal alpha is obtained by averaging across 15 bootstrapped held-out sets. In addition to fitting models with WordNet features extracted 4s prior to the measured neural response, we developed longer timescale linear models by concatenating the WordNet features extracted for each second (as described above) over T-second windows with T ranging from 1 to 20 seconds and presented these aggregated features to the bootstrapped regularized regression model. Figure S4 (B) demonstrates the performance of WordNet models across different groups of regions as a function of T, and (C) depicts the voxel-level prediction accuracy (R) of the best performing WordNet model that stacks features from 4-12s (at an interval of 1s) prior to the encoded cortical response. While simple and interpretable, the WordNet models clearly under-perform in terms of prediction accuracy (R) in comparison to the models proposed in the present study.

**Fig. S3.**
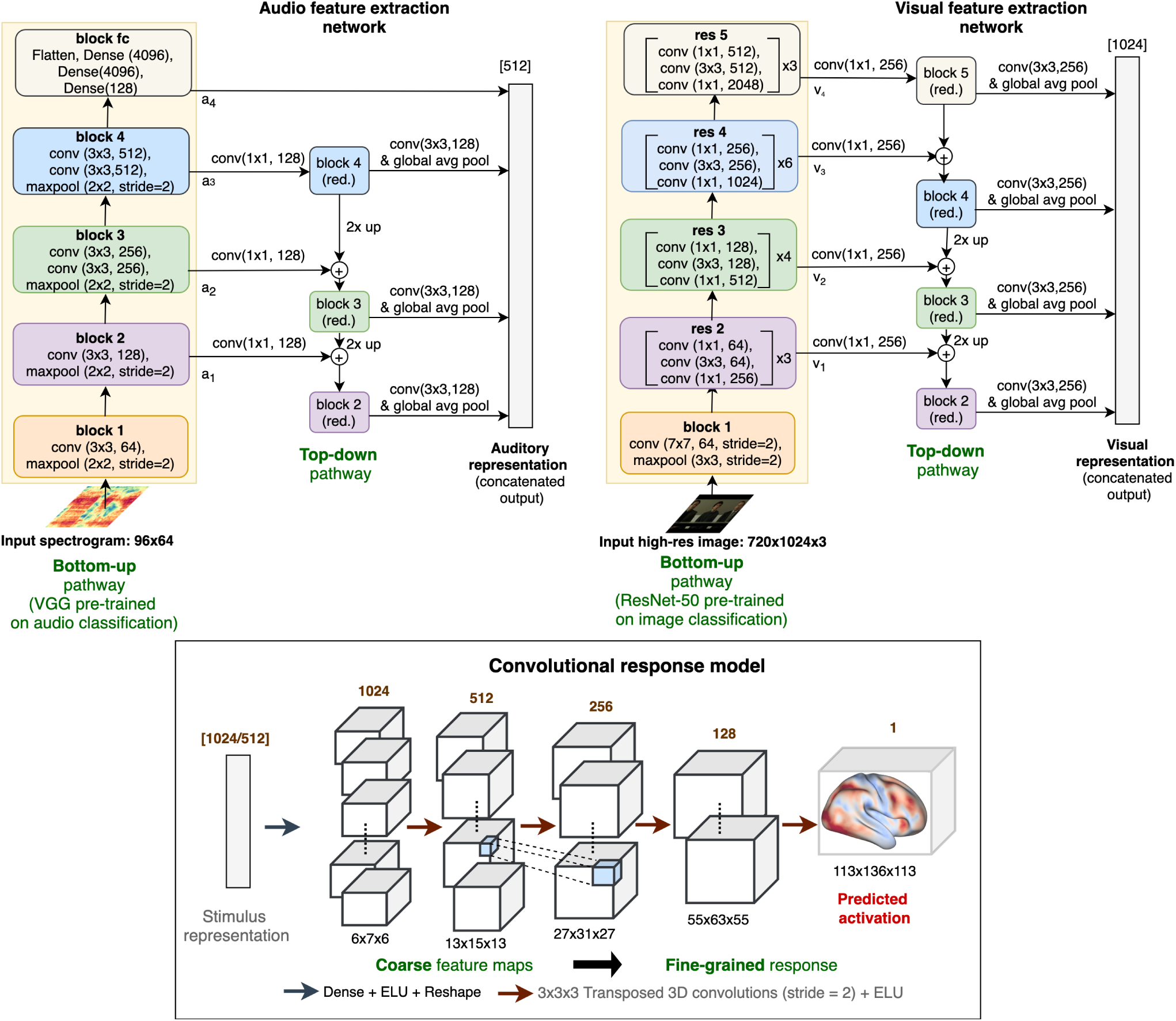
Implementation details for the audio (top left) and visual (top right) feature extraction networks as well as the convolutional response model (bottom). All layers and blocks outside the yellow rectangle (bottom-up pathway) are trained from scratch. The blocks inside the yellow rectangular window are initialized with networks pre-trained on image or sound recognition. Further, ResNet-50 is frozen during the training of all encoding models, whereas VGG is fine-tuned. The sequence of operations within each block are defined from top to bottom, while the number of repetitions for each sequence within the block are indicated with the multiplicative symbol on the right.

### Regularized linear regression: deep convolutional features

We also trained group-level encoding models using a linear response model since this constitutes the dominant state-of-the-art approach to neural encoding (*5, 4, 7*). To enable a fair comparison against the proposed 1-sec uni-modal models, we extract hierarchical features from the same layers of the ResNet-50 and VGG-ish architectures as employed by the proposed models. The only difference here is the lack of a top-down pathway (since it is not a part of the pre-trained network but is trained with random initialization on the neural response prediction task), which prevents the refinement of coarse feature maps before aggregation. Pooling the outputs of different layers channel-wise using the global average pooling operation (namely {*v*_1_, *v*_2_, *v*_3_, *v*_4_} for the visual model and {*a*_1_, *a*_2_, *a*_3_, *a*_4_} for the audio model in Figure S3) leaves us with and 1024 and 3840 features to present to the auditory and visual models, respectively. Further, to compare against the longer-duration 20-sec models, we adopted two approaches: (1) we simply concatenated the stimulus features extracted for each second (as described above) over T-second windows with T ranging from 1 to 20 seconds and presented these aggregated features to the linear response model; alternatively, (2) we reduced the dimensionality of the aggregated features to a fixed length (set to 128) as in (1) using principal component analysis run on the training data. We added this comparison to rule out the fact that the temporal trend in performance of linear models is simply driven by a higher-dimensional feature space. We note that even after dimensionality reduction, the components retained at least 80% of the explained variance in all cases. Audio-visual encodings with linear response models were obtained similarly by simply fusing the respective audio and visual hierarchical features through concatenation before linear regression. We apply *l*_2_ regularization on the regression coefficients and adjust the optimal strength of this penalty through cross-validation on the training data using log-spaced values in {1e –14, 1e14} for each model. We report performance of the best models in Figure S4(A). Note that unlike the WordNet models, we found that optimizing a single regularization penalty *α* common across all voxels outperformed independent voxelwise fitting with bootstrap in this case. Thus, we only present the results for the former. We note here that the convolutional response model in our proposed approach (instead of a fullyconnected approach) allowed us to keep the learnable parameters manageable, facilitating joint optimization/fine-tuning of the feature extractor and response models. The consistently superior performance of the proposed models against linear regression based approaches strongly suggests that there is merit in end-to-end learning for encoding responses to dynamic, multi-sensory stimuli.

**Fig. S4.**
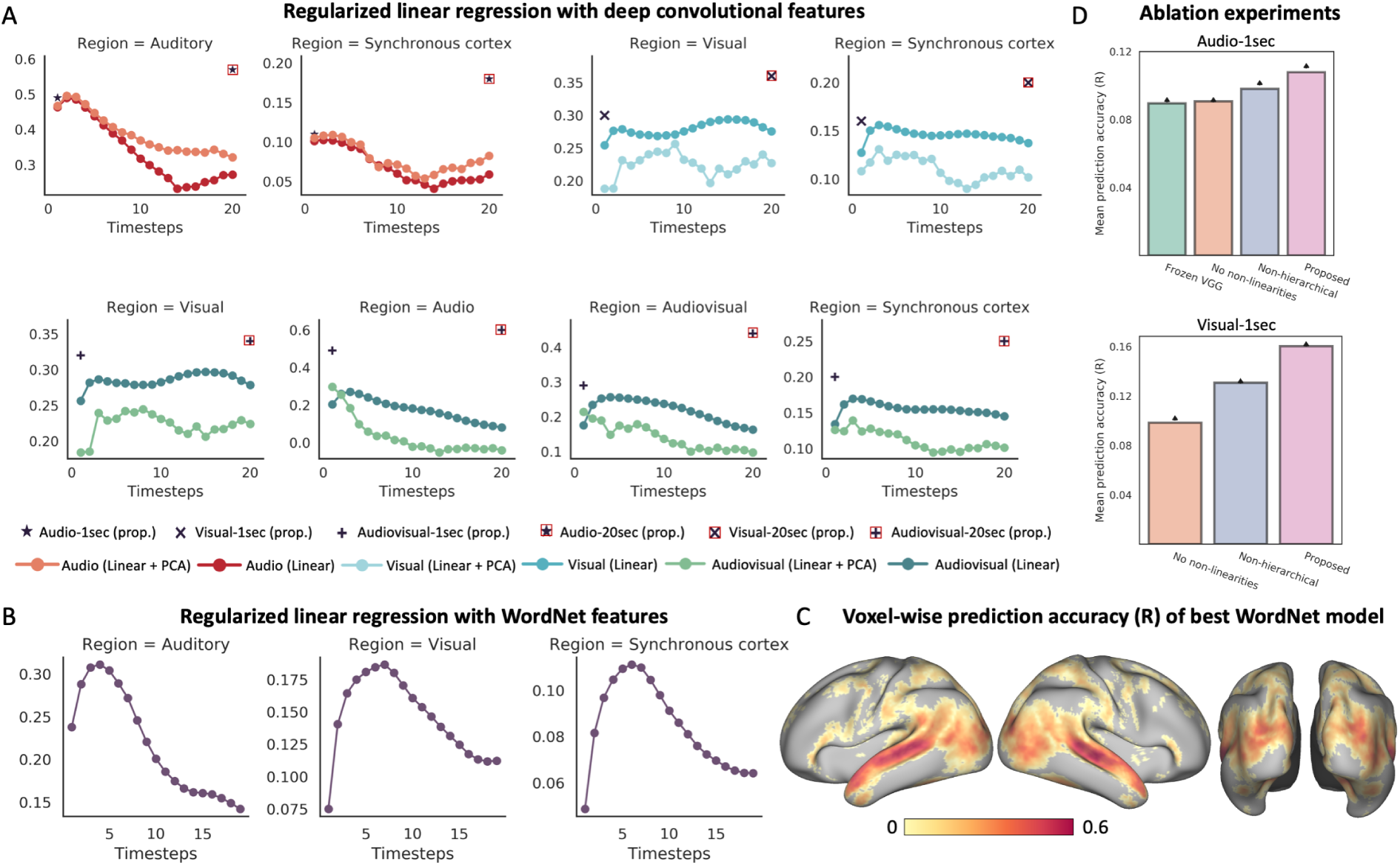
Performance of linear response models with (A) deep convolutional features and (B) semantically rich WordNet features. The x-axis depicts the length of the windows (in seconds) over which the stimulus features are concatenated and y-axis shows the mean Pearson’s correlation coefficient between the predicted and measured responses across the stimulus-driven voxels. (C) shows the cortical map of the prediction accuracy (R) for the best WordNet model. (D) shows results of the ablation study and highlights the importance of different components of the proposed model architecture.

### Ablation study

To determine the influence of different architectural components on prediction performance of the proposed models, we performed an ablation study to investigate the individual contributions of (i) non-linearities in the response model, (ii) hierarchical (multi-scale) feature maps and (iii) fine-tuning audio sub-network (VGG). We selectivity removed each of these components from the respective 1sec models and compared the resulting performance against the proposed model that employs all (i)-(iii) components. There are several interesting observations to make from this ablation analysis (Figure S4D). (i) First, we find that encoding models with a frozen VGG network that is not updated during training incur a loss in performance compared to the proposed model where VGG layers are trainable during neural response prediction. This clearly demonstrates the advantages of altering these pre-trained models and suggests that fine-tuning is both feasible and beneficial in improving neural response prediction. (ii) Next, we find that prediction performance deteriorates after removing the non-linearities in both the Audio-1sec and Visual-1sec models. In the context of the Visual-1sec model with a frozen pre-trained backbone (ResNet-50) and coupled with (i), this observation further highlights that it is possible to develop models of human sensory processing that are quantitatively more precise in matching brain activity than task-driven neural networks. (iii) Finally, we assessed the benefit of using hierarchical feature maps over selecting the single best-performing layer for each model (audio or visual) based on cross-validation. For both audio and visual models, we find that features from the last layer (i.e., *a*_4_ and *v*_4_, respectively) yield the highest mean prediction accuracy (R) across the synchronous cortex. However, although the convolutional response model architecture is common across these encoding models, it is important to note that this analysis is still plagued by confounds such as the different dimensionality of feature spaces across different layers that feed into the response model. The best performing single-layer encoding model, however, still performs worse than the hierarchical approach.

### Computing significance estimates

The statistical significance of individual voxel predictions (Figure 3) was computed as the pvalue of the obtained sample *correlation coefficient* for the null hypothesis of uncorrelation (i.e., true correlation coefficient is zero) under the assumptions of a bivariate normal distribution. We employed the false-discovery procedure of Benjamini & Hochberg (1995) (*51*) to control for multiple comparisons under assumptions of dependence. For statistical comparison of model performance within each group of regions in Figure 2 (main text), we performed paired t-test on ROI-level average performance metrics and corrected for multiple comparisons among models (Bonferroni).

### Sensory-sensitivity index

Distorting the input to the audio-visual model at test time allows us to interrogate the sensory-sensitivity of different brain regions. We developed a sensory-sensitivity index of each ROI based upon predictive performance of the model with distorted inputs, as shown in Figure 5. Let *SV*_*r*_ and *SA*_*r*_ denote the mean prediction accuracy of the model in region *r* after shuffling (temporally) the input order of the visual and auditory stimuli, respectively. The sensory-sensitivity index for region *r* is then defined as 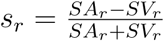. Note that positive values of this index indicate that region *r* incurs a greater loss in predictivity upon distortion of visual information than auditory information, suggesting a higher visual sensitivity for this voxel. Similarly, negative values signal towards a higher auditory-sensitivity.

### Stimuli for synthetic contrasts

Synthetic contrasts were generated to study the generalization of our models to new experimental paradigms (Figure 6). We focus on predicting task-based contrasts for three semantic categories, namely, *faces, places* and *speech*, since these are the most well-studied categories in the context of their distinct functional signatures. The stimuli for visual contrasts were derived from the HCP Working Memory paradigm, which combines category specific representation tasks (including faces and places) and working memory tasks. After excluding gray-scale images, we were left with 102, 77, 97 and 103 images for the categories of faces, places, body parts and tools, respectively. Since these are static image without any dynamic content, we employed the Visual-1sec model to derive the visual contrasts (Figure 6(C),(D)).

Stimuli for the speech and non-speech contrast were extracted from large popular datasets for these categories. Speech stimuli were extracted from a human speech-utterance dataset comprising short audio clips of interviews recorded on YouTube (*52*). Non-speech stimuli were extracted from another large dataset comprising short clips of environmental sounds (*53*). We randomly extracted ∼100 minutes of audio waveforms from these datasets for both categories. The stimuli were processed for mel-spectrogram extraction in the same manner as the HCP audio-visual movies. Since the non-speech stimuli only comprised contiguous clips of roughly 3 – 5 second duration, we employed the Audio-1sec model to obtain the speech contrast (Figure 6(B)).

### Performance improvement and autocorrelation decay

In the past, processing timescales in the brain have been probed using several different means (*46*). In one of the proposed approaches, the decay time of temporal autocorrelation is used as a proxy measure to understand the variation in processing timescales across different brain regions. With this approach, it was shown that decay times increased progressively along the temporal hierarchy. Following this line of work, we estimated the autocorrelation decay time constant (*π*) for each voxel by fitting an exponential, *A* exp {–*t/π*}, to the autocorrelation function (autocorrelation computed at different lags). The exponential model was first independently fit for each movie run and each voxel and the estimated *π* were subsequently averaged across runs to obtain one decay time constant per voxel. Here, we were primarily interested in understanding whether there is any relationship between the performance improvement of the 20-sec model over 1-sec model, Δ*R*, computed as the difference between the prediction accuracies of the Audiovisual-20sec and Audiovisual-1sec at every voxel, and the temporal autocorrelation properties of that voxel. We hypothesized that in voxels with longer processing timescales, the autocorrelation would persist for longer durations (resulting in larger *π*) and the longer timescale model (20-sec) would yield more substantive improvement over the 1-sec model. As shown in Figure S5, we observed a significantly positive correlation between performance improvement and the autocorrelation decay time constant (r = 0.49 and 0.50 across voxels in auditory and visual regions as defined in Table S2), in line with our hypothesis. This suggests that the benefit of employing the 20-sec model, as quantified in terms of performance improvement, is indeed more remarkable in regions with longer processing timescales.

**Fig. S5.**
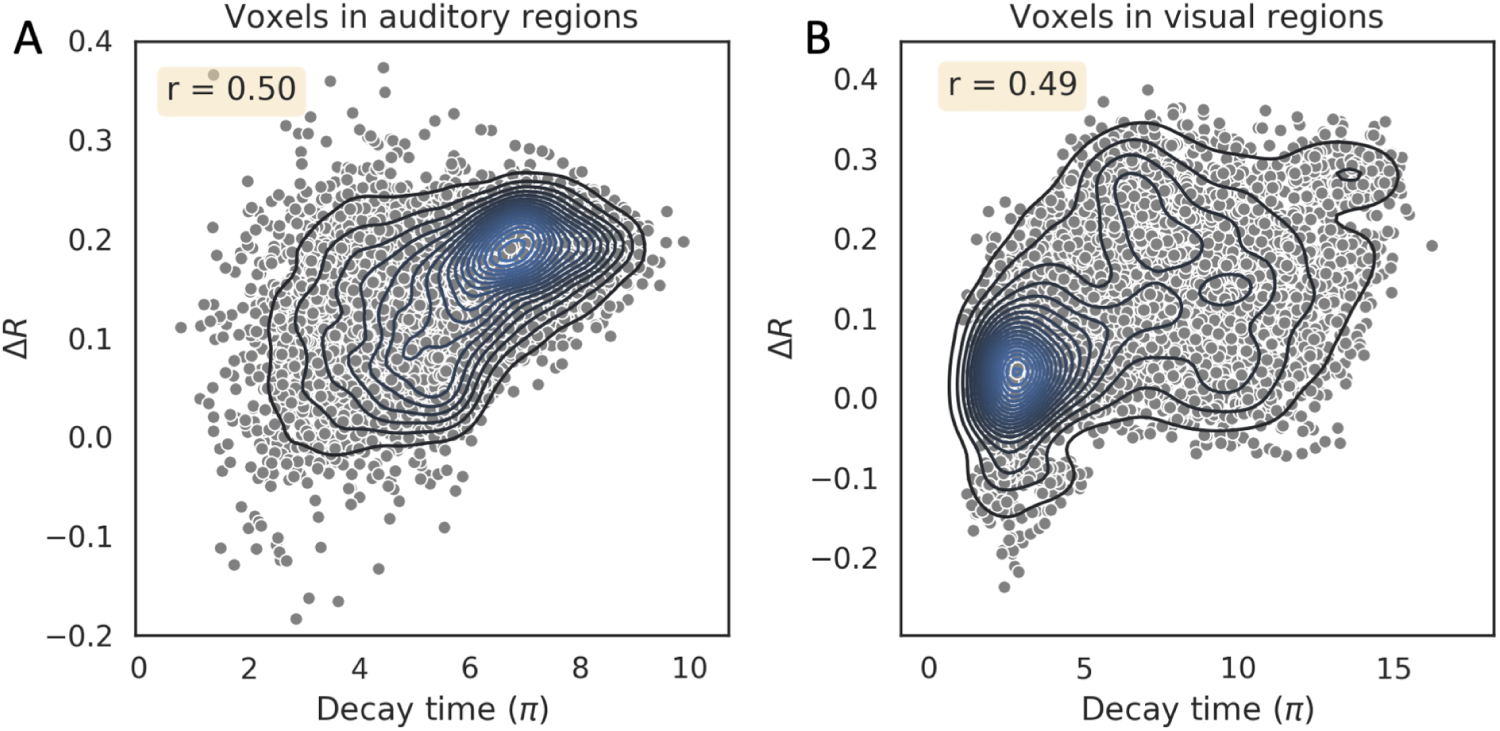
Performance boost of the 20-sec model over 1-sec model is higher in voxels with longer autocorrelation decay times. (A) & (B) depict the performance improvement (Δ*R*) against decay time constants for voxels associated with auditory and visual regions, respectively (Table S2). The *r* value indicates the Pearson’s correlation coefficient between the two quantities. Each dot in the scatterplot represents an individual voxel. Bivariate kernel density estimates are overlaid on top of the scatterplot as contours to depict the probability distribution of observations.

### Surface visualization

All input fMRI data, as well as response predictions in this study are volume based. In order to be consistent with prior research on encoding models that employ surface visualizations, we created surface versions of volumetric predictability and synthetic contrast maps, as shown in Figures 3, 5 and 6. We employed the 3D trilinear mapping method from connectome workbench that computes the result on each vertex based on linear interpolation from voxels on each side of the vertexfc2 ^2^. However, since volume to surface mappings are an approximation, we only employ this conversion for visualizations. All reported metrics are computed on volumes only on a per-voxel basis.

### Qualitative analysis

To gain qualitative insights into the predictions of the most accurate model (Audiovisual-20sec) on the held-out movie, we plot the predicted as well as measured response time-series of the voxel with ‘median’ prediction accuracy (R) in the best performing ROI of each group (Figure S6). The latter corresponds to A4, V3CD, STSdp, IFSp and Area 45 for the auditory, visual, multi-sensory, frontal and language groups respectively.

**Fig. S6.**
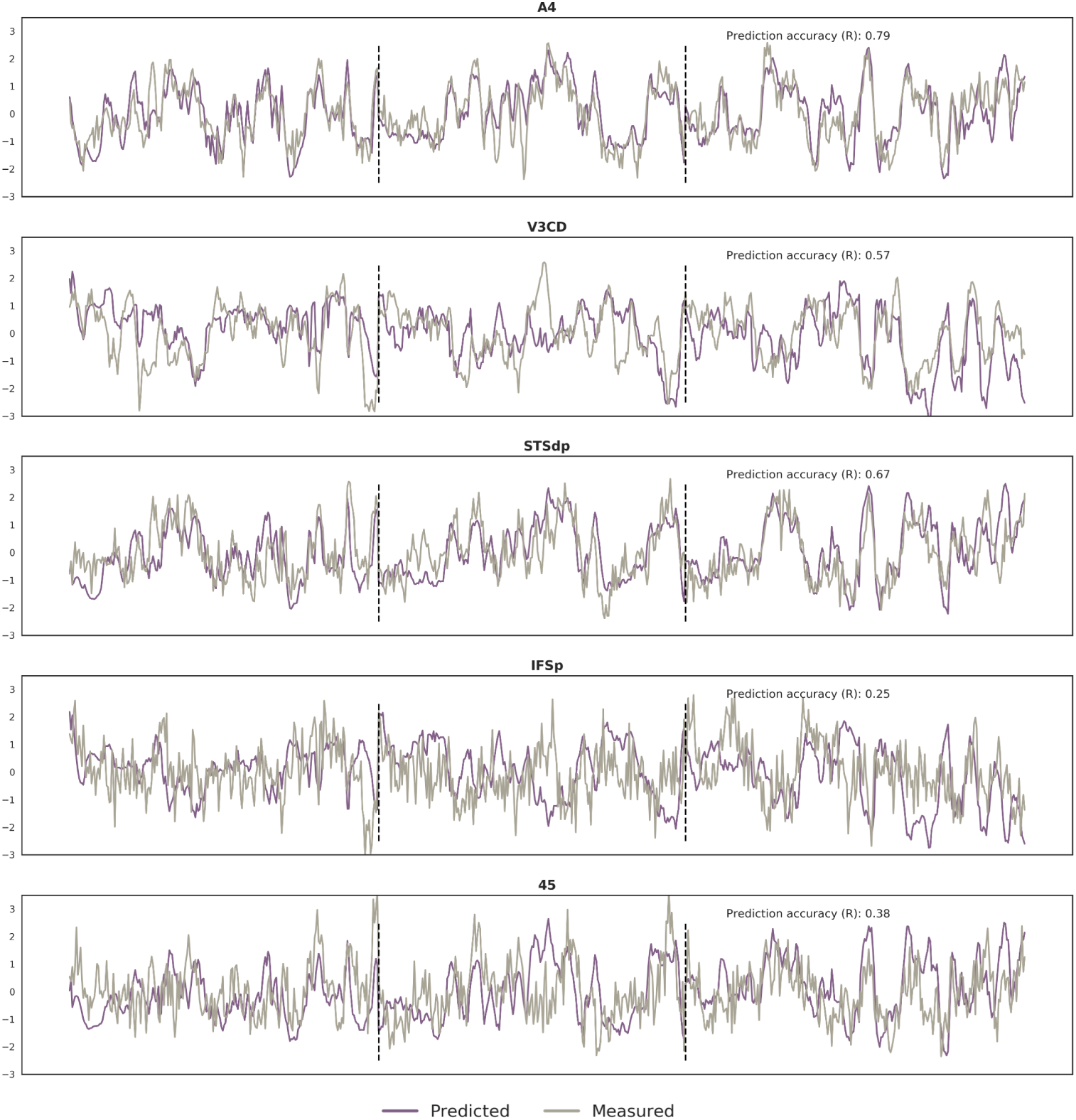
Predicted and measured response time-series of the ‘median’ predictive accuracy (R) voxel across ROIs of different functional groups. Vertical dashed lines mark the boundary of clip segments in the held-out movie.

### Group-level prediction accuracy: held-out set

To test the generality of the models, we further compared model predictions against the group-averaged response of a held-out group within HCP comprising 20 novel subjects distinct from the 158 individuals used in the training set, on the same independent held-out movie.

#### Noise ceiling estimation

For the held-out group, we obtain the noise ceiling by considering variability across subjects. Here, the noise ceiling was computed as the correlation coefficient between the mean measured response for the *independent* test movie across all 158 subjects in the training set and the group-averaged response computed over the 20 new subjects. This metric captures the response component shared across independent groups of subjects and thus reflects the upper bound achievable by a group-level encoding model. We employ this noise ceiling for comparison against the prediction accuracy of the model on the held-out group of subjects (Figure S7).

The models accurately predicted cortical responses evoked by the *independent* test movie as measured in the *independent* subject population (Figure S7, S8), with the best performing model (Audiovisual-20sec) even achieving close to perfect predictivity relative to the “noise ceiling” in certain multi-sensory sites such as the posterior STS (Figure S7(A), (G)). Here, the noise ceiling was computed as the correlation coefficient between the mean neural response in the *independent* test movie, across all 158 subjects in the training set and the group-averaged response computed over the 20 new subjects. This metric captures the response component shared across independent subject populations and thus reflects the upper bound achievable by a group-level encoding model. These results clearly indicate that inclusion of temporal history and multi-sensory information pushes the prediction accuracies closer to their upper bound, as also evidenced by a higher slope of the linear model fit on their corresponding data points. Further, voxels that truly approach the noise ceiling are predominantly associated with the auditory group of regions as broadly characterized within the HCP MMP parcellation. Interestingly, we find that this regional distribution of predictivity against noise ceiling holds even for subject-specific responses and not just the group-averaged responses, as described in the next section and shown in Figure S9.

### Subject-level prediction accuracy: held-out set

For each participant in our independent subject group (N = 20), we computed the correlation coefficient (R) between the predictions of the best performing model (Audiovisual-20sec) and the subject-specific fMRI response corresponding to the independent movie. We further contrast this cortical map of prediction performance against another map computed as the voxel-wise correlation coefficient between the mean neural response across all 158 training subjects and the respective subject-specific response on the independent movie. The latter places an upper bound on the predictivity of each voxel as achievable by any group-level model. Here, we present the results for 5 subjects with mean prediction accuracy (un-normalized) within the stimulus-driven cortex in the *i*th percentile with *i ∈ {*0.01, 25, 50, 75, 99.9*}*. The results (Figure S9) suggest that the model can successfully capture the response component that individual subjects share with the population.

**Fig. S7.**
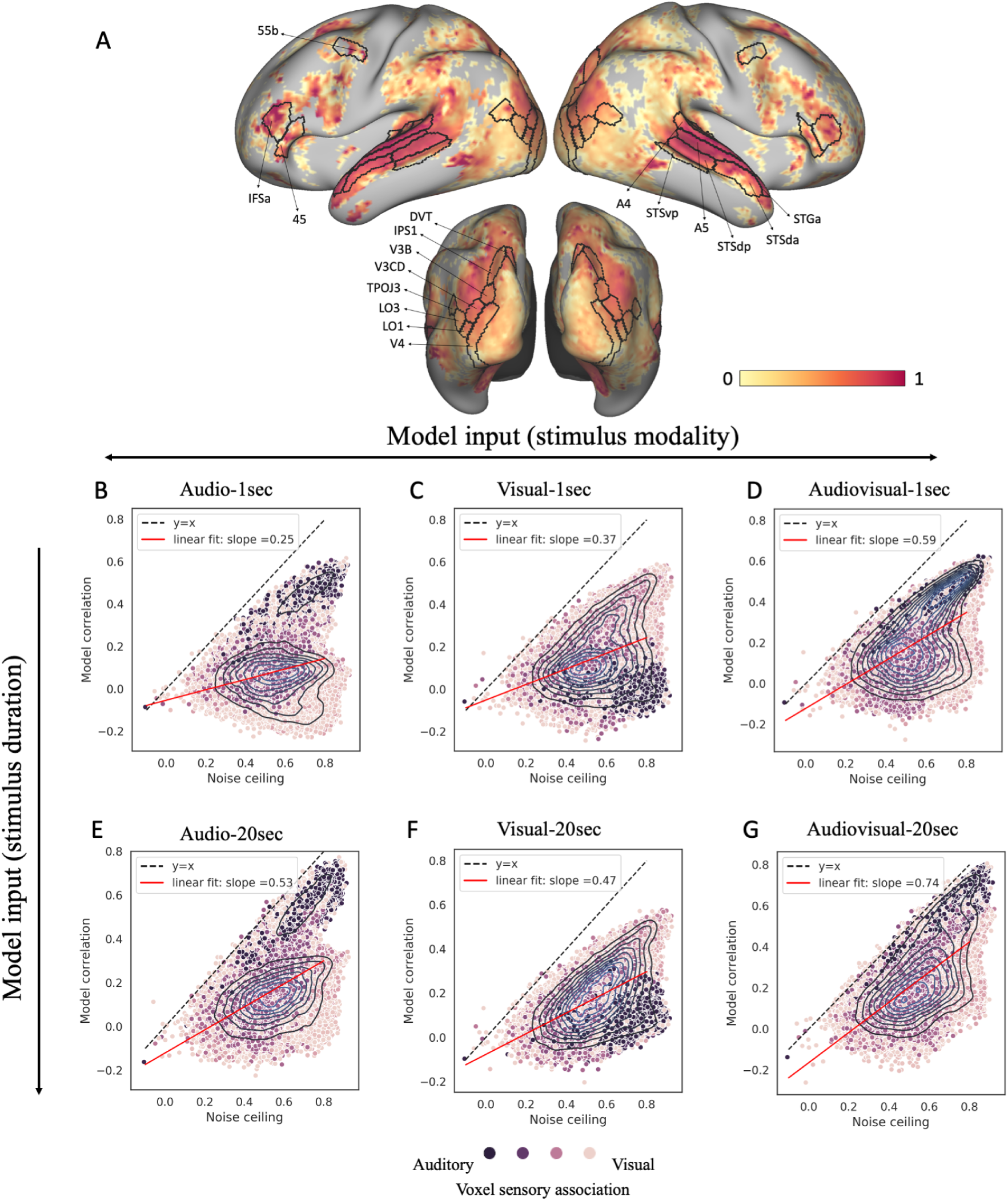
Model performance on held-out group of subjects. (A) Pearson’s correlation coefficient (R) between the model predictions and group-averaged response of an independent subject group comprising 20 subjects, on the held-out test movie, normalized by the voxel-specific noise ceiling. (B) Predictivity against the noise ceiling for all voxels with high “synchrony” across training movies (*>*0.5) (see Supplementary Information for details). This gives a total of 52,954 highly “synchronous” voxels that are colored based on their association with auditory and visual groups. This hue assignment of each voxel was derived from the coloration of the corresponding ROI in the multi-modal HCP parcellation. Each dot in the scatterplot represents an individual voxel. Bivariate kernel density estimates are overlaid on top of the scatterplot as contours to depict the probability distribution of observations (prediction accuracy/noise ceiling pair at every voxel).

**Fig. S8.**
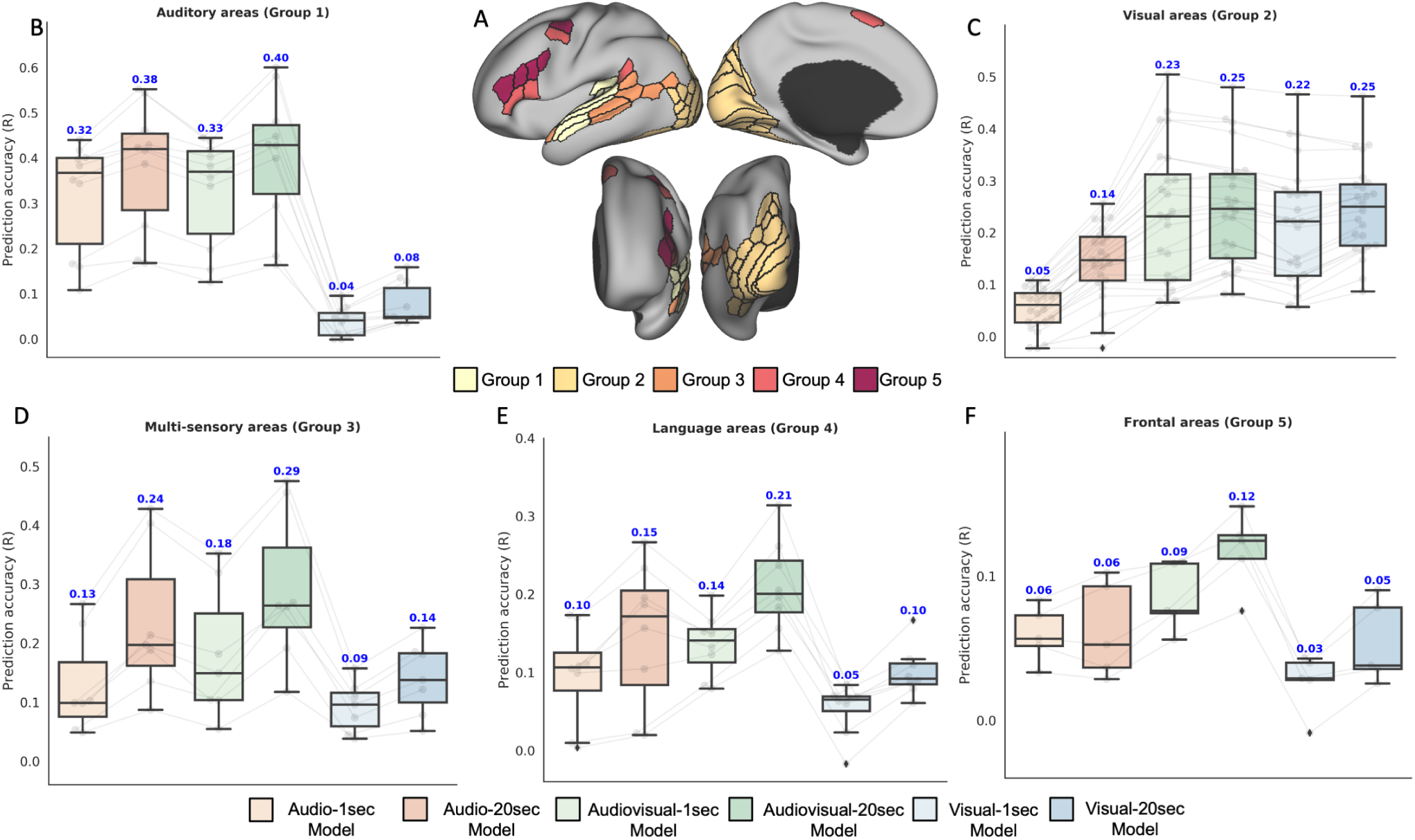
Quantitative evaluation metrics for all the proposed models on the independent *held-out* population comprising 20 novel subjects. (B)-(F) depict prediction accuracy (R) for all the proposed models across major groups of regions as identified in the HCP MMP parcellation (A). Predictive accuracy of all models is summarized across (B) auditory, (C) visual, (D) multi-sensory, (E) language and (F) frontal areas. Box plots depict quartiles and swarmplots depict mean prediction accuracy of every ROI in the group. For language areas (Group 4), left and right hemisphere ROIs are shown as separate points in the swarmplot because of marked differences in the prediction accuracy. Statistical significance tests (results indicated with horizontal bars) are performed to compare 1-sec and 20-sec models of the same modality (3 comparisons) or uni-modal against multi-modal models of the same duration (4 comparisons) using paired t-test (p-value *<* 0.05, Bonferroni corrected) on mean prediction accuracy within ROIs of each group.

**Fig. S9.**
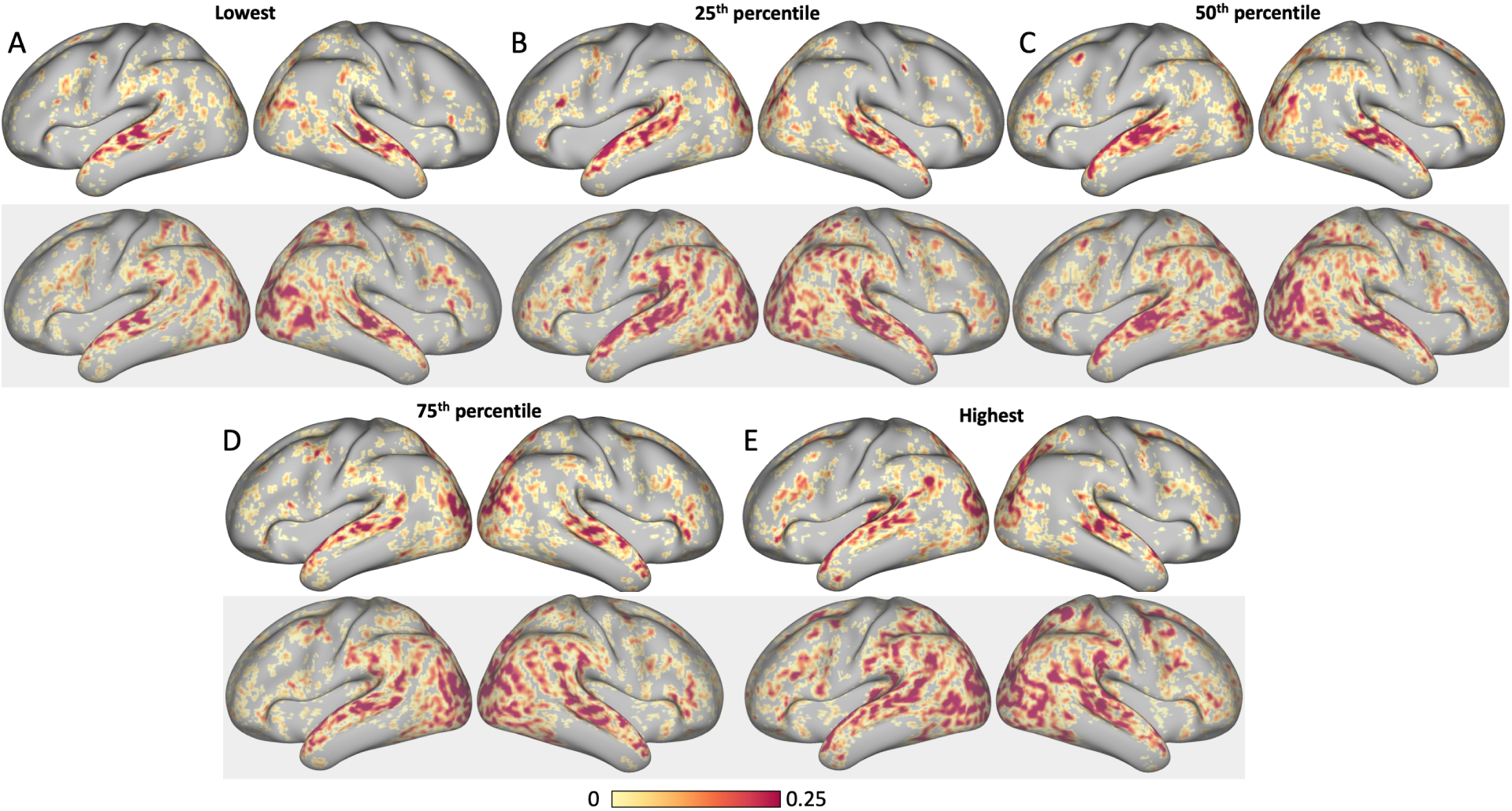
Comparison of voxel-level prediction accuracies (R) against subject-specific noise ceiling for 5 representative subjects from the held-out set. The subjects were chosen such that their mean prediction accuracy (unnormalized) within the stimulus-driven cortex lied in the *i*th percentile with *i ∈* {0.01, 25, 50, 75, 99.9}. Surface maps with white background in (A)-(E) depict raw correlation coefficients between model (Audiovisual-20sec) predictions and subject-specific response on the held-out movie whereas maps on gray background indicate the respective subject-specific noise ceiling. Only significantly correlated voxels (p*<*0.05, FDR corrected) are colored on the surface.

## Footnotes

1 Pre-trained tensorflow/keras models for the visual and auditory backbone were available at https://keras.io/applications https://github.com/tensorflow/models/tree/master/research/audioset/vggish-respectively

2 https://www.humanconnectome.org/software/workbench-command

